# Genomic stability of Self-inactivating Rabies

**DOI:** 10.1101/2020.09.19.304683

**Authors:** Ernesto Ciabatti, Ana González-Rueda, Daniel de Malmazet, Hassal Lee, Fabio Morgese, Marco Tripodi

**Affiliations:** MRC Laboratory of Molecular Biology, Cambridge, United Kingdom

## Abstract

Transsynaptic viral vectors provide means to gain genetic access to neurons based on synaptic connectivity and are essential tools for the dissection of neural circuit function. Among them, the retrograde monosynaptic ΔG-Rabies has been widely used in neuroscience research. A recently developed engineered version of the ΔG-Rabies, the non-toxic self-inactivating (SiR) virus, represents the first tool for open-ended genetic manipulation of neural circuits. However, the high mutational rate of the rabies virus poses a risk that mutations targeting the key genetic regulatory element in the SiR genome could emerge and revert it to a canonical ΔG-Rabies. Such revertant mutations have recently been identified in a SiR batch. To address the origin, incidence and relevance of these mutations, we investigated the genomic stability of SiR *in vitro* and *in vivo*. We found that “revertant” mutations are rare and accumulate only when SiR is extensively amplified *in vitro*, particularly in suboptimal production cell lines that have insufficient levels of TEV protease activity. Moreover, we confirmed that SiR-CRE, unlike canonical ΔG-Rab-CRE or revertant-SiR-CRE, is non-toxic and that revertant mutations do not emerge *in vivo* during long-term experiments.

**Highlights:**

- Revertant mutations are rare and do not accumulate when SiR is produced in high-TEVp expressing production cell lines
- SiR is non-toxic *in vivo*
- Revertant SiR mutations do not accumulate during *in vivo* experiments

## Introduction

The development of innovative technologies to record and manipulate the activity of large populations of neurons (Jun et al., 2017; Lin and Schnitzer, 2016; Stirman et al., 2016; Yizhar et al., 2011) has had a transformative impact on systems neuroscience leading to a deeper understanding of how specific networks control essential aspects of animal behaviour (Fadok et al., 2017; Kohl et al., 2018; Stuber and Wise, 2016). In particular, the latest generation of molecular sensors and actuators allow researchers to visualize (Abdelfattah et al., 2019; Dana et al., 2019) and perturb (Kato et al., 2018; Shemesh et al., 2017) the activity of individual neurons with unprecedented genetic, spatial and temporal resolution. However, strategies to express these tools in any desired neuron within a neural network structure remain scarce. Viral vectors represent the primary approach to deliver genetic materials to mammalian brains, with adeno associated viruses (AAV) rapidly becoming the primary choice to target neurons based on anatomical location, genetic identity or projection pattern (Chan et al., 2017; Tenenbaum et al., 2004; Tervo et al., 2016). Nonetheless, transsynaptic viruses are the only vectors that are able to label cells based on their synaptic connectivity, permitting the functional dissection of neural circuits. Among them, the retrograde monosynaptic G-deleted Rabies virus (ΔG-Rabies) is the most sensitive and efficient transsynaptic retrograde tracer, widely used to highlight the structural organization of neural networks in mammals (Callaway and Luo, 2015; Stepien et al., 2010; Tripodi et al., 2011; Wickersham et al., 2007b). However, its toxicity has limited its use for functional experiments. Indeed, in the past few years, several strategies have been applied trying to overcome the known toxicity of rabies vectors and extending their use for long-term functional interrogation of neural circuits: the use of different viral strains (CVS-N2c) (Reardon et al., 2016), the conditional destabilization of viral proteins (Self-inactivating Rabies, SiR)(Ciabatti et al., 2017) or the deletion of essential genes other than G (ΔGL-Rabies)(Chatterjee et al., 2018).

All these approaches have advantages and disadvantages and collectively represent important improvements in the Rabies design. For example, the use of different parental strains in ΔG-Rabies vectors provide delayed mortality and improved tropism (Reardon et al., 2016), but do not overcome the continuous viral replication that eventually leads to toxicity. The deletion of genes other than G gave origin to effective axonal retrograde tracers (Chatterjee et al., 2018) but requires the expression of multiple transgenes for transsynaptic tracing experiments via other viruses or using transgenic animals, which have yet to be fully implemented and that risk recreating a fully functional ΔG-Rabies in the starter cells. The addition of regulatory elements to the rabies genome, as in the SiR design in which the rabies nucleoprotein (N) is conditionally targeted to the proteasome by a PEST sequence, has the advantage of abolishing continuous viral replication (Ciabatti et al., 2017). On the other hand, the known high mutation rate of RNA viruses (Drake and Holland, 1999; Sanjuan et al., 2010) poses the risk that naturally occurring mutations could emerge to selectively inactivate the added genetic sequence, hence potentially giving origin to toxic revertant mutants.

In its original design, SiR is produced from cDNA in conditions where PEST is constantly removed by the tobacco etch virus protease (TEVp) cleavage, which should prevent accumulations of PEST-targeting mutations. While it was suggested that such PEST-targeting mutations might be an unavoidable outcome of the SiR design (Chatterjee et al., 2018), here we show that such mutations, in fact, only accumulate when SiR is extensively amplified in cells expressing suboptimal levels of TEVp. Conversely, minimizing the number of passages *in vitro* and using high-TEVp expressing production cell lines prevents any appreciable accumulation of such mutations during SiR production.

The reported findings that ΔG-Rabies-CRE showed an apparently reduced cytotoxicity (Chatterjee et al., 2018) led to the suggestion that the CRE expression alone could dampen the toxicity of all ΔG-Rabies vectors, and hence of the SiR-CRE as well (Matsuyama et al., 2019). However, the survival of a fraction of ΔG-Rabies-CRE-infected neurons in CRE-reporter mice might be explained by the presence of a few naturally occurring defective viral particles that lack one or more key viral genes (Wiktor et al., 1977), which could effectively recapitulate the self-inactivating behaviour purposefully engineered in the SiR virus. Indeed, here we show that CRE expression alone is ineffective in dampening toxicity and that while SiR-CRE is entirely non-cytotoxic in cortical and sub-cortical regions for several months, canonical ΔG-Rabies-CRE displays a significant toxicity *in vivo*.

In summary, here we investigated the genomic stability of SiR and found that when produced in cells with high levels of TEVp with few rounds of amplification PEST-targeting mutations do not accumulate to appreciable levels. As expected, revertant-free SiR-CRE viruses but not Rab-CRE or PEST-mutated SiR-CRE are entirely non-toxic. Moreover, we show that PEST-targeting mutations do not accumulate at appreciable rate *in vivo*.

## Results

### *De novo* SiR productions do not accumulate revertant mutations

SiR self-inactivation depends on the proteasomal targeting of N by the c-terminal addition of a PEST sequence. The high rate of mutation in RNA viruses (10^−6^ to 10^−4^ substitutions per nucleotide per round of copying) (Sanjuan et al., 2010) could lead to the emergence of mutations targeting PEST. If these mutations generate a premature stop codon just upstream of the c-terminal PEST sequence they could effectively revert the SiR to a canonical and cytotoxic ΔG-Rabies. To address the issue of whether and/or to what extent the emergence of such “revertant” mutants occurs we generated 8 independent SiR productions from cDNA following the protocol we previously described (Ciabatti et al., 2017). We produced viral genomic libraries for each preparation (50 clones/batch) for Sanger sequencing using primers carrying random octamers in order to identify individual particles (Fig. 1A-B). Out of the 8 independent preparations for a total of 400 individually analysed particles, we did not identify particles harbouring the non-sense mutations described by Matsuyama and colleagues (Fig. 1B and Table 1). The sequences’ analyses showed the presence of sporadic mutations across other genomic locations (Table 1) as expected given the rabies mutational rate. Notably, several clones per preparation had point mutations within the N/P intergenic region, suggesting that the stop-polyadenylation signal is permissive to single base mutations (Table 1). These data confirm that SiRs generated from cDNA as described in Ciabatti et al. (2017) do not accumulate mutations upstream the PEST domain at appreciable levels.

**Fig 1.**
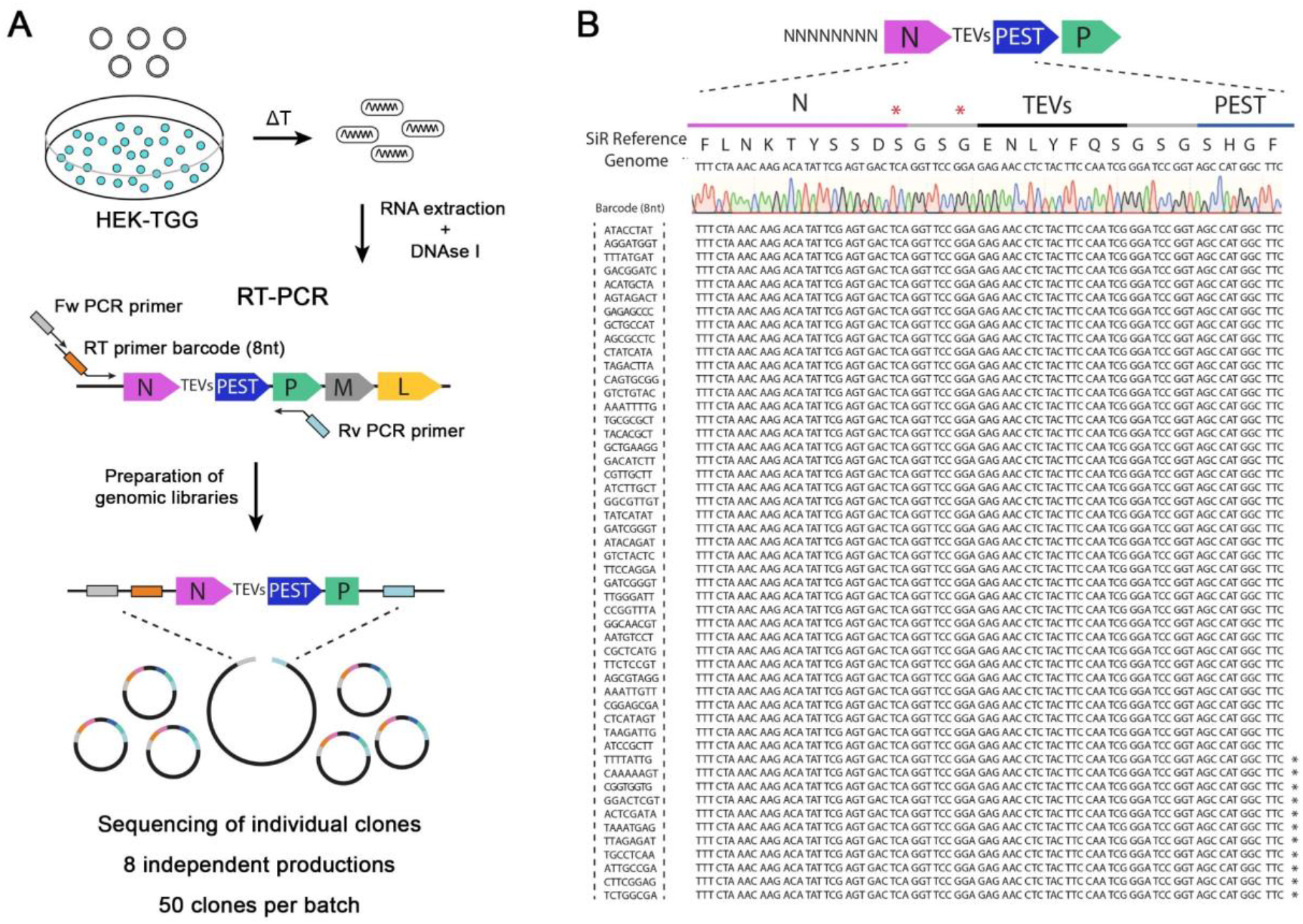
SiR production from cDNA leads to revertant-free viral preparations. A) Scheme of experimental strategy to identify the emergence of “revertant” mutations during SiR production. 8 independent SiR preparations were rescued from cDNA and genomic RNA were extracted, treated with DNAse I, subjected to RT-PCR to amplify N-TEVs-PEST coding sequence and used to generate libraries for Sanger sequencing (50 clones per preparation were sequenced). B) Example of sequencing results from one SiR preparation showing no mutations at the end of N. Stars (*) show the position of previously identified mutations, marks on the sequences indicates the presence of mutations in different positions.

### Analysis of molecular mechanisms underpinning the potential emergence of SiR revertant mutants

Although we found no indication of emergence of PEST-targeting mutations when SiR is rescued from cDNA, a recent report finding two batches of PEST-mutated SiR (Matsuyama et al., 2019) unarguably points to the possibility of emergence of these mutations under certain conditions. Hence, we sought to determine which conditions might favour the accumulation of revertant mutants. In the SiR design, the PEST sequence is fused to the N protein through a cleavable linker that allows its efficient production from TEVp-expressing packaging cells (Ciabatti et al., 2017). The constant removal of PEST ensures that naturally occurring mutations that inactivate PEST do not provide advantage over non-mutated particles. However, we reasoned that with suboptimal TEVp activity PEST-mutants may display faster replication kinetics than SiR particles, and might eventually accumulate in the population, as in a directed-evolution experiment. Thus, we hypothesised that two factors might prominently affect the emergence of revertants: 1. low TEVp levels in packaging cells and 2. excessive rounds of amplification of SiR *in vitro*. First, we investigated TEVp activity in packaging cells over time by producing HEK293T cells expressing TEVp and Gsad (HEK-TGG) as previously described (Ciabatti et al., 2017). After selecting for TEVp-expressing cells with puromycin HEK-TGG where cultured for multiple passages in medium containing different level of antibiotic (puromycin 0 μg/ml, 1 μg/ml, 2 μg/ml; Fig 2A). TEVp activity was then assessed every 2 passages by transfecting a TEVp reporter (Gray et al., 2010) and analysing TEVp site (TEVs) cleavage by western blot (Fig 2B, Fig S1). We found that the TEVp-dependent cleavage of the overexpressed reporter decreased in HEK-TGG after amplification and by passage 6 (P6) was less than half the initial level (from 31.7 ± 2.4% at P0 to 14.7 ± 1.7% and 13.8 ± 1.2% with 1 μg/ul and 2μg/ul puromycin, respectively; Fig 2B-C). Importantly, amplification in the absence of antibiotic pressure quickly reduced TEVp activity, decreasing by one order of magnitude by P6 (31.7 ± 2.4% at P0; 7.7 ± 1.3% at P2; 3.1 ± 0.2% at P6 without puromycin; Fig 2B-C). This suggests that extensive amplification of HEK-TGG leads to selection of clones with suboptimal TEVp expression, particularly in absence of antibiotic pressure.

**Fig 2.**
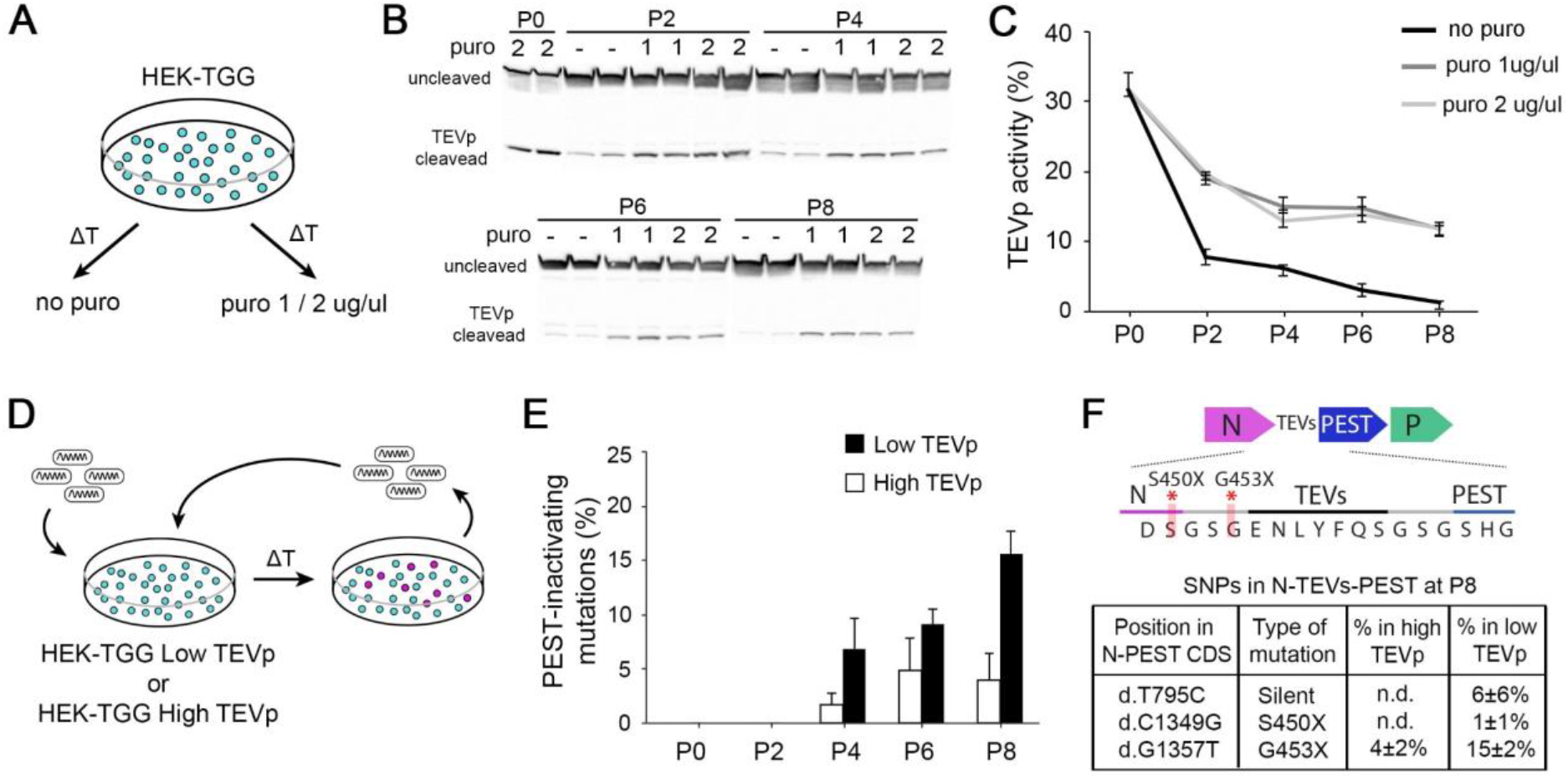
High TEVp activity in packaging cells prevents accumulation of PEST-mutations. A) HEK-TGG packaging cells were amplified for several passages in absence or presence (1 or 2 μg/ml) of puromycin selection. B) TEVp-dependent cleavage of TEVp-activity reporter was analysed by western blot in HEK-TGG at different amplification passages. C) Quantification of TEVp-activity in packaging cells over time in presence or absence of antibiotic pressure. (mean ± SEM, n = 3) D) Experimental design to assess emergence of mutations in SiR preparations after multiple passages of amplification in high TEVp (HEK-TGG P0) or low TEVp HEK-TGG (HEK-TGG P8, without puromycin selection). E) Quantification of frequency of the accumulation of PEST-targeting mutations over time that prevent translation of PEST domain. (mean ± SEM, n = 4 independent viral preparation). F) Summary of the single nucleotide polymorphisms (SNPs) in the coding sequence (CDS) of N-TEVs-PEST that reached threshold at P8 (mean ± SEM, n = 4) (n.d. indicates that the mutations were not detected above threshold). Top scheme shows the position of PEST-inactivating mutations.

To test the dependence of the emergence of revertant mutations on TEVp activity in the packaging cells, and investigate the accumulation kinetics of potential mutants we amplified 4 independent (sequenced) revertant-free SiR preparations *in vitro* in low- and high-TEVp conditions for several passages. Every two passages, genomic libraries for each viral preparation were produced by retrotranscribing the RNA genomes using primers barcoded with unique molecular identifiers (UMI, random decamer) and PCR amplifying an amplicon containing the N-TEVs-PEST gene. Then, SiR libraries were analysed by long-read next generation sequencing (NGS) using single molecule, real-time (SMRT) PacBio technology (Rhoads and Au, 2015) (Fig 2D and Fig S2). SMRT sequencing employs the generation of circular molecules from the N-TEVs-PEST amplicons that are replicated for several passages by a polymerase so that individual sub-reads can be combined to generate high quality consensus sequences (sequencing accuracy ≥98% with 3 passages; Fig S2). Since SMRT technology is particularly prone to false-positive insertion and deletions (INDELs)(Carneiro et al., 2012; Dohm et al., 2020) and all previously reported PEST-targeting mutations were substitutions (Matsuyama et al., 2019), we restricted our analysis to substitutions (single nucleotide polymorphism, SNP) above 2% threshold. We considered a PEST-targeting mutation to be any non-synonymous substitution targeting either N or TEVs-PEST sequences. In accordance with our hypothesis, the extensive amplification of SiR *in vitro* led to the emergence of revertants that can accumulate within the SiR population, especially in low-TEVp packaging cells (16%± 2% of sequences containing a revertant mutation at P8 in low-TEVp cells; FiG 2E, Table 2). On the other hand, PEST-targeting mutations remained below 5% even after 8 rounds of amplification when SiR was amplified in high-TEVp cells (4%± 2% of sequences containing a revertant mutation at P8 in high-TEVp cells; Fig 2E, Table 2). Notably, all PEST-inactivating mutations detected in this experiment were single base substitutions introducing a premature stop codon prior to TEVs either at the last amino acid of N or immediately after (d.C1349G and d.G1357T, leading to stop insertion at S450 and G453, respectively; Fig 2F, Table 2), which also accounted for the large majority of revertant particles reported by Matsuyama et al. (2019). Thus, in order to avoid the accumulation of revertant mutants, SiR viruses should be only amplified in high-TEVp, low-passage packaging cells for the minimum required number of passages.

### Difference in cytotoxicity between ΔG-Rabies, PEST-mutant SiR and SiR

In the recent report of Matsuyama et al. (2019) the authors showed that PEST-mutant SiR is cytotoxic *in vivo*, which is the obvious consequence of the presence of a stop codon upstream PEST that transforms the SiR into a WT ΔG-Rabies. This is strikingly different to our results showing that SiR can permanently label neurons by recombinase-mediated activation of genetic cassettes before disappearing from the infected neurons without cytotoxicity (Ciabatti et al., 2017). To experimentally confirm that revertant-free and PEST-mutant SiR are different viruses we characterized them *in vitro* and *in vivo* and compared them to canonical ΔG-Rabies. In order to obtain a pure preparation of PEST-mutants we engineered each of the two nonsense mutations previously reported (Matsuyama et al., 2019) (d.C1349G and d.G1357T, leading to stop insertion at S450 and G453, respectively; Fig 2F) in the SiR cDNA, generating two viruses named SiR-S450X and SiR-G453X (Fig 3A and S3). First, we confirmed the loss of functional TEVs in the PEST linker in the engineered-revertants by observing the TEVp-dependent virally driven GFP expression *in vitro* (Fig S3). Next, we assessed the *in vivo* cytotoxicity of SiR, SiR-G453X and ΔG-Rab expressing CRE by injecting them in the CA1 hippocampal region of CRE-dependent tdTomato reporter mice and analysing the number of infected neurons at different time points post injection (p.i.) as in our previous study (Ciabatti et al., 2017)(Fig 3B). In line with our published results no mortality was detected in SiR-infected hippocampi (4109 ± 266 tdTomato+ neurons at 1 week p.i.; 4458 ± 739 tdTomato+ neurons at 2 months p.i.; one-way ANOVA, F = 0.08, p = 0.92, Fig 3C-D) while only 44% of tdTomato+ neurons were detected in Rabies-targeted and 60% in SiR-G453X-targeted hippocampi at 2 months p.i. (1422 ± 184 at 1 week versus 624 ± 114 at 2 months p.i. for ΔG-Rab; one-way ANOVA, F = 11.55, p = 0.003; 3052+508 at 1 week versus 1829+198 at 2 months p.i. for SiR-G453X; one-way ANOVA, F = 4.27, p = 0.05; Fig 3C-D). These results confirm the requirement for an intact PEST sequence to sustain the self-inactivating behaviour of SiR and suggest that PEST-targeting mutations do not occur *in vivo*. Interestingly, a fraction of tdTomato+ neurons survived in ΔG-Rab-CRE–injected brains, differing from what we observed when injecting ΔG-Rab-GFP, where no cells were detected at 3 weeks p.i. (Fig 3CD) (Ciabatti et al., 2017). To experimentally confirm that revertant particles indeed do not emerge *in vivo* during long-term SiR experiments, we prepared NGS libraries of SiR genomes extracted from hippocampi of injected animals before SiR switch off and sequenced them by SMRT sequencing (Fig 3E and S2). In all 3 independent experiments no revertant mutations had accumulated *in vivo* above threshold prior to the switching off of the virus (Fig 3F, Table 3).

**Fig 3.**
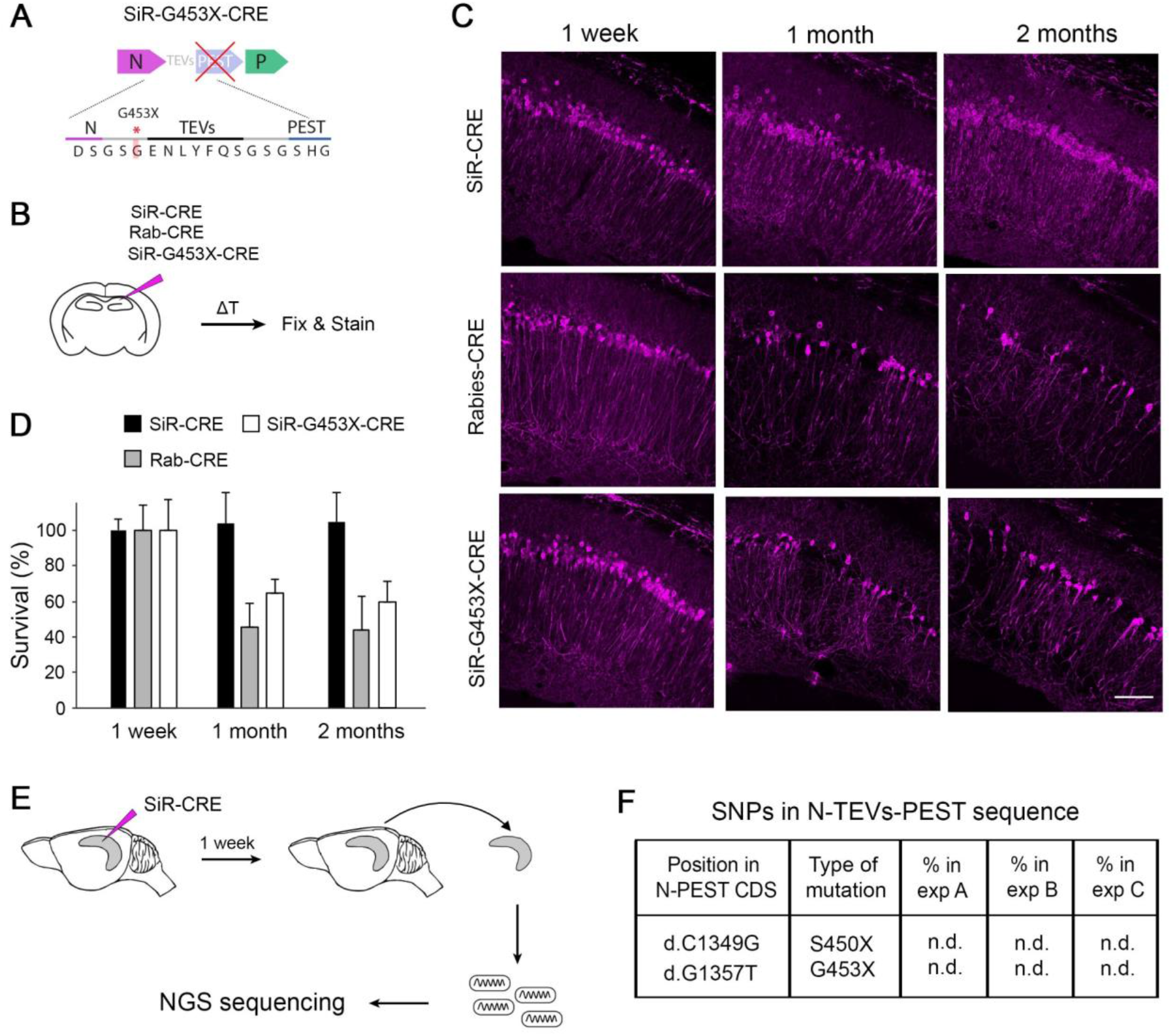
Revertant-free SiR, but not PEST-mutant, is non-toxic and does not accumulate PEST-targeting mutations *in vivo*. A) Scheme of the engineered PEST-mutant SiR (SiR-G453X). B) Experimental procedure. C) Confocal images of hippocampal sections of *Rosa-LoxP-STOP-LoxP-tdTomato* mice infected with SiR-CRE, Rab-CRE, SiR-G453X and imaged at 1 week, 1 month and 2 months p.i. Scale bar, 50 μm. D) Number of tdTomato positive neurons at 1 week, 1 months, and 2 months p.i. normalized to 1-week time point (mean ± SEM, n = 4 animals per virus per time point). E) Experimental procedure for the sequencing of SiR particles from injected hippocampi at 1 week p.i. F) List of PEST-inactivating mutations above 2% thresholds with relative frequency in each animal (n.d. indicates that the mutation was not detected above threshold). (n = 3 animals).

To further confirm the lack of any toxic effect in SiR-targeted neurons we also performed longitudinal imaging of cortical neurons using 2-photon microscopy. These longitudinal experiments allowed us to follow the morphology and survival of the same identified SiR-targeted neurons over time in living mice, thereby giving more direct evidence of the potential cytotoxicity or lack thereof associated with SiR. We imaged SiR-CRE or ΔG-Rab-CRE labelled neurons in the cerebral cortex of *Rosa-LoxP-STOP-LoxP-tdTomato* mice for up to 5 months p.i. (Fig 4A-B). The total number of detectable tdTomato^+^ neurons increased in SiR injected animals between 1 and 2 weeks and remained constant for the entire duration of the experiment (Fig 4B), while ΔG-Rab-injected cortices show a decrease of total number of tdTomato^+^ neurons over time (Fig 4B). Importantly, nearly all the SiR-targeted neurons imaged at 1 week were detected in subsequent imaging sessions (97% ± 1 tdTomato^+^ at 21 weeks p.i.; Fig 4C) in contrast to ΔG-Rab-infected neurons, where ~70% of the neurons detected at 1 week had died by 9 weeks p.i. (29% ± 2 tdTomato^+^ at 21 weeks; Fig 4C). These results show virtually no loss of SiR labelled neurons during the entire imaging period (five months) and confirm the lack of any observable cytotoxic effect of SiR on the recipient neurons (Fig 4B-D).

**Fig 4.**
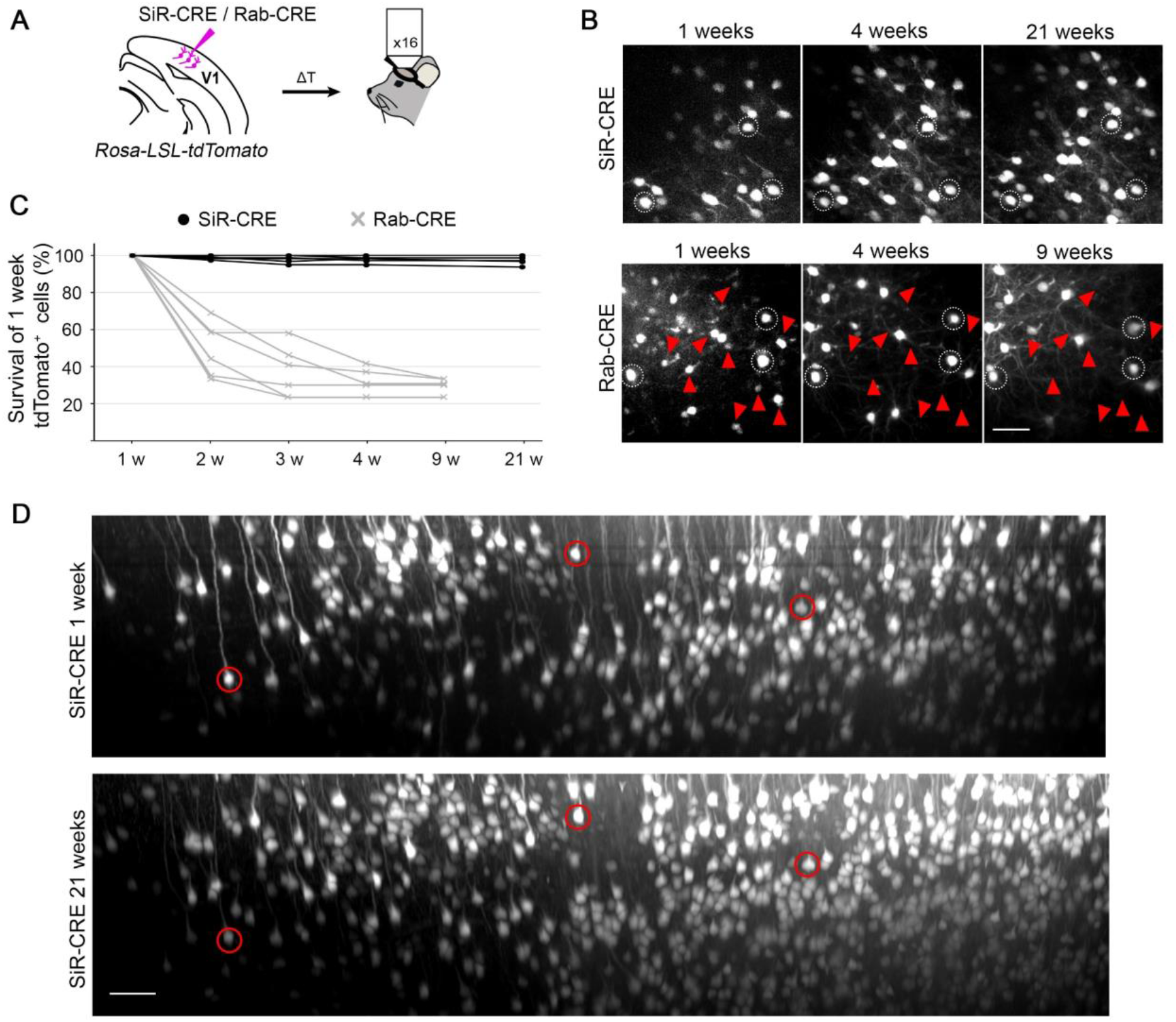
2-photon in vivo longitudinal imaging of revertant-free SiR-infected cortical neurons reveals no toxicity and unaltered neuronal morphology after 5 months. A) Schematic of SiR-CRE or Rab-CRE injection in *Rosa-LoxP-STOP-LoxP-tdTomato* mice in V1 followed by *in vivo* imaging. B) 2-photon maximal projection of the same field in SiR-CRE and Rab-CRE injected cortices at 1, 4 and 21 weeks p.i. or 1,4 and 9 weeks, respectively. Red arrowheads mark tdTomato positive neurons detected at 1 week that disappear in later recordings. Scale bar 50 μm. C) Survival of the tdTomato positive cells recorded at 1 week over time. (ROIs = 6 per virus. n = 2 animals per virus). D) 2-photon maximal projection of the same large field in SiR-CRE injected cortices at 1 week and 21 weeks p.i. Scale bar 50 μm.

## Discussion

The development of technologies to record and perturb the activity of neurons within neural circuits has been instrumental for the recent progress in systems neuroscience. ΔG-Rabies viruses have been transformative in the study of neural circuit organization in animal models, especially mammals. The recent generation of a non-toxic SiR vector has opened the door to the long-term functional dissection of neural networks. One concern regarding its widespread use has been the risk that mutations could emerge and compromise SiR preparations by reverting the SiR vector to canonical and cytotoxic ΔG-Rabies.

Here we have investigated the genomic stability of SiR and showed that PEST-targeting mutations are rare and do not accumulate when SiR is produced directly from cDNA as previously described. However, we show that revertant mutants can emerge if SiR is extensively amplified *in vitro*, particularly in cells expressing suboptimal levels of TEVp, where revertant mutants have a specific replication advantage. Nonetheless, we also show that when production utilises HEK-TGG packaging cells expressing high levels of TEVp, even 8 rounds of amplification *in vitro* do not lead to the accumulation of PEST-targeting mutations above 5%. Notably, we found that TEVp activity inevitably decreases after several passages of amplification of HEK-TTG, thus fresh low passage packaging cells should always be used to produce SiR preparations.

Another important question is, when revertant-free SiR is produced and used for tracing experiments, can PEST-targeting mutations emerge *in vivo*? Here we show that revertant-free SiR-CRE efficiently traces neurons *in vivo* without toxicity in cortical and subcortical regions for several months p.i.. Importantly, PEST-mutant SiR is as toxic as canonical ΔG-Rabies, indicating that an intact PEST sequence is essential for SiR non-toxic behaviour and suggesting that revertant mutants do not emerge during *in vivo* experiments. We confirmed this by sequencing the SiR viral particles isolated from *in vivo* experiments and found no PEST-targeting mutations. Thus, the short lifetime of the SiR in the infected neurons does not permit PEST-mutations to emerge and accumulate *in vivo* before viral disappearance when revertant-free SiR preparations are used.

Although PEST-inactivating mutations can be prevented during production and do not accumulate *in vivo*, strategies to further reduce or entirely eliminate the risk of their appearance could simplify viral production in other laboratories and allow the use of SiR in sensitive applications, e.g. re-targeting the same starter cells multiple times. In our experiments only two specific revertant mutations were identified, single base substitutions that introduce a stop signal either at the last amino acid of N or in the linker prior to TEVs and PEST (d.C1349G and d.G1357T) which accounted for the large majority of revertant mutations found in Matsuyama et al. (2019). Future studies should focus on investigating if this and other potential hotspots in the SiR genome can be optimised to simplify the production of SiR.

## Supporting information

Reference sequence for NGS data

## Author Contributions

M.T. and E.C. conceived the project. E.C. designed the experiments. E.C. performed all the experiments with the exception of the 2-photon recordings. F.M. provided technical support for cell culture and molecular biology. A.G.-R performed mouse surgeries. D.D.M. performed mouse surgeries and the *in vivo* 2-photon imaging experiments. H.L performed molecular biology experiments. E.C. and D.D.M. analysed the data. E.C. prepared the figures. E.C. wrote the manuscript incorporating feedback from M.T., A.G.-R., D.D.M. and H.L.

## Acknowledgments

We would like to thank Elena Williams for comments on the manuscript. We thank Jerome Boulanger for writing the script for the 3d-alignment of 2-photon recordings, Nicolas Alexandre for the help with the bioinformatic analysis of the NGS datasets, the Laboratory of Molecular Biology (LMB) workshops for the help with software and hardware development, and members of the Biological Service Group for their support with the *in vivo* work. This study was supported by the Medical Research Council (MC_UP_1201/2), the European Research Council (STG 677029 to M.T.). All data are stored on the LMB server. All materials described in this paper can be obtained upon reasonable request and for non-commercial purposes after signing a material transfer agreement (MTA) with the MRC.

## Methods

### Contact for Reagents and Resource Sharing

Further information and requests for resources and reagents should be directed to the corresponding author: Ernesto Ciabatti.

### Experimental Model and Subject Details

#### Animal strains

C57BL/6 wild type (WT) mice and *Rosa-LoxP-STOP-LoxP-tdTomato* transgenic mice (Jackson: Gt(ROSA)26Sortm14(CAG tdTomato) were used. All procedures were conducted in accordance with the UK Animals (Scientific procedures) Act 1986 and European Community Council Directive on Animal Care. Animals were housed in a 12 hours light/dark cycle with food and water ad libitum.

#### Cell lines

HEK293T cells were obtained from ATTC. HEK293T packaging cells expressing Rabies glycoprotein (HEK-GG) were generated by lentivirus infection with Lenti-H2BGFP-2A-GlySAD and after 3 passages GFP expressing cells were selected by fluorescent activated cell sorting (FACS). HEK293T packaging cells expressing Rabies glycoprotein and TEV protease (HEK-TGG) were generated from HEK-GG by lentivirus infection with Lenti-puro-2A-TEV and selected, after 3 passages, with 1 μg/ml of puromycin added to the media for 1 week.

### Method Details

#### Design and generation of ΔG-Rabies and SiR plasmids

All Rabies and SiR plasmids were generated by Gibson cloning starting from pSAD-ΔG-F3 plasmid (Osakada et al., 2011) or SiR vectors we previously generated (Ciabatti et al., 2017), respectively. Engineered SiR vectors carrying d.C1349G or d.G1357T PEST-targeting mutations were produced by PCR amplification of the Rabies genome in 2 fragments starting from the end of N assembled using Gibson master mix (NEB).

The lentiviral vectors used to generate the packaging cells have been previously described (Ciabatti et al 2017).

#### TEVp activity in packaging cells

Low passage HEK-TGG packaging cells were produced as previously described (Ciabatti, 2017). Briefly, HEK293T cells were infected with Lenti-GFP-2A-G and after 3 passages GFP expressing cells were selected by fluorescent activated cell sorting (FACS). Cells were infected with Lenti-puro-2A-TEVp and amplified for 2 passages under 2 μg/ml of puromycin selection in 10% DMEM. This produced the HEK-TGG P0 line that was further amplified either in absence or presence of 1/2 μg/ml of puromycin selection for up to 8 passages. Cells were split every 3 days at 1:6 dilution and every 2 passages TEVp activity was assessed by seeding 750k cells in 6-wells and transfecting a TEVp activity reporter (Gray et al., 2010) after 24 hrs. Transfected cells were lysed in RIPA buffer after 24 hrs and TEVp-dependent reporter cleavage was assessed by western blot.

#### Viral productions

SiR and ΔG-Rabies viruses were rescued from cDNA by the co-transfection of rabies genome vectors with pcDNA-T7, pcDNA-SADB19N, pcDNA-SADB19P, pcDNA-SADB19L, and pcDNA-SADB19G (Osakada et al., 2011) in HEK-TGG and HEK-GG cells, respectively, as previously described (Ciabatti et al., 2017).

For the recovery of high titer SiR and ΔG-Rabies, HEK-TGG or HEK-GG respectively were infected in 15 cm dishes at ~80% confluence with 3 ml of viral supernatant obtained as described in the viral screening section. Cells were split the day after infection and maintained for 1 or 2 days at 37°C and 5% CO_2_ checking daily the viral spreading when a fluorescent marker was present. Then, the media was replaced with 2% FBS DMEM and maintained for 2 days at 35°C and 3% CO_2_. Viral supernatant was collected, cell debris removed by centrifugation at 2500 rpm for 10 minutes followed by filtration with 0.45 μm filter and the virus concentrated by ultracentrifugation on a sucrose cushion as previously described (Wickersham et al., 2007a).

#### Ontogenesis of revertant mutations during viral production

8 independent SiR viruses were rescued from cDNA as described in previous section. SiR RNA genomes were extracted from the infectious supernatants with RNeasy kit (Qiagen) following manufacturer’s instructions and used to generate plasmid libraries for Sanger sequencing. To investigate the emergence of mutations during subsequent viral amplification rounds *in vitro* low passage HEK-TGG (HEKTGG P0), or high passage cells amplified in absence of puromycin pressure (HEK-TGG P8) were seeded in 10 cm dishes. At 60-70% confluence cells were infected with SiR supernatants obtained from cDNA at MOI= ~2-3. The next day cells were split at 1:2 dilution and maintained for 1 day at 37°C and 5% CO_2_ in 10% FBS DMEM. Then, media was replaced with 2% FBS DMEM and cells moved to incubation at 35°C and 3% CO_2_. Viral supernatants were collected after 2-3 days and used to infect fresh HEKTGG P0 or HEK-TGG P8. The entire process was repeated for a total of 8 rounds of viral amplification. At each passage 1 ml of supernatant was used to extract viral RNA genomes and generate libraries for NGS.

#### Analysis of SiR accumulation of mutations during in vivo experiments

Sequence-verified revertant-free SiR virus was injected in CA1 region of the hippocampus of C57BL/6 wild type mice. After 1 week, mice were culled and the injected hippocampi manually dissected immediately. SiR genomes were obtained by homogenising the hippocampi with Tissuelyser II (Qiagen) and extracting the total RNA with RNeasy kit (Qiagen) according to manufacturer instructions. 500 ng of RNA per hippocampus were retrotranscribed using superscript IV kit (Invitrogen) and amplicons of N-TEVs-PEST were PCR-amplified to generate libraries for SMRT NGS sequencing.

#### Sanger sequencing of SiR genomes

SiR genomic copies were extracted by concentrating 1 ml of infectious supernatant with Amicon Ultra-4 10K filters in an Eppendorf 5810R centrifuge at 4C, 2500g for 20’ followed by RNeasy kit (Qiagen) extraction. RNA samples were treated with DNAse I (Invitrogen) for 15’ at RT followed by inactivation at 65C for 10’. Genomes were retro-transcribed with SuperScript IV Reverse Transcriptase (Invitrogen) following manufacturer instructions using a primer complementary to the 5’ leader sequence containing an 8 nt random barcode:

Leader_8barcode_:

TCAGACGATGCGTCATGCNNNNNNNNACGCTTAACAACCAGATC

cDNA samples were subjected to RNAse H treatment (NEB) followed by PCR amplification of a fragment corresponding to the entire coding sequence of N-TEVs-PEST and part of the P gene with Platinum SuperFi II Master Mix polymerase (denaturation for 30 s at 98C; 25 cycles of amplification with 5 s at 98C, 10 s at 60C and 60 s at 72C; 3 min at 72 for final extension) using primers:

Leader_PCR_Fw: ccaccgcggtggcggccgctcTCAGACGATGCGTCATGC

P_PCR_Rv: ctaaagggaacaaaagctgggtacCTTCTTGAGCTCTCGGCCAG

The obtained ~2Kb amplicons were gel purified from 1% agarose gel using QIAquick Gel Extraction Kit (Qiagen) and cloned in pBluescript SK (+) (GenBank:X52325.1) digested KpnI - XbaI using Gibson assembly cloning method (NEB). 50 clones were purified and sequenced by Sanger method using M13_Fw and M13_Rv primers checking that each sequence carried a different 8 nt barcode.

#### Single molecule real-time (SMRT) sequencing of SiR genomes

SiR supernatant preparations were first concentrated by centrifuging 1 ml of infectious supernatant in Amicon Ultra-4 10K filters in an Eppendorf 5810R centrifuge at 4C, 2500g for 20’, followed by RNA extraction using RNeasy kit (Qiagen). Purified viruses were directly extracted with RNeasy kit by adding 350 ul of RT lysis buffer to 5 ul of concentrated virus. RNA samples were treated with DNAse I (Invitrogen) for 15’ at RT followed by inactivation at 65C for 10’. Genomes were retro-transcribed with SuperScript IV Reverse Transcriptase (Invitrogen) following manufacturer instructions using a primer complementary to the 5’ leader sequence containing an adapter sequence and a 10 nt random barcode:

Pacbio_Leader_10barcode:CGAACATGTAGCTGACTCAGGTCACNNNNNNNNNNCACGCTTAACAACCAGATC

cDNA samples were subjected to RNAse H treatment (NEB) followed by PCR amplification of a fragment corresponding to the entire coding sequence of N-TEVs-PEST and a fragment of the P gene with Platinum SuperFi II Master Mix polymerase (denaturation for 30 s at 98C; 25 cycles of amplification with 5 s at 98C, 10 s at 60C and 60 s at 72C; 3 min at 72 for final extension) using primers asymmetrically barcoded as shown below (list of the barcodes used for each sample can be found in Tables 2,3):

Pacbio_PCR_Fw: (16nt_barcode)CGAACATGTAGCTGACTCAGGTCAC

Pacbio_PCR_Rv: (16nt_barcode)AGTCGCCCCATATCCTCAGG

Barcodes:

bc1: TCAGACGATGCGTCAT

bc2: CTGCGTGCTCTACGAC

bc3: CATAGCGACTATCGTG

bc4: GCTCGACTGTGAGAGA

bc5: ACTCTCGCTCTGTAGA

bc6: TGCTCGCAGTATCACA

bc7: CAGTGAGAGCGCGATA

bc8: TCACACTCTAGAGCGA

bc9: GCAGACTCTCACACGC

bc10: GTGTGAGATATATATC

bc11: GACAGCATCTGCGCTC

bc12: CTGCGCAGTACGTGCA

The obtained ~2Kb amplicons were gel purified from 1% agarose gel using QIAquick Gel Extraction Kit (Qiagen) followed by clean-up with QIAquick PCR purification kit (Qiagen). Purified barcoded amplicons from different viral preparations were combined in a single tube to obtain equimolar ratio and final concentration of ~50 ng/ul. SMRTbell libraries of pooled amplicons (up to 29 samples per library) were prepared using SMRTbell Template Prep Kit 1.0 (PAcbio) and Sequel chemistry v3 and sequenced on a PacBio Sequel SMRT cell with a 10 hr. movie.

#### Single molecule real-time (SMRT) sequencing analysis

Pacbio Sequel II raw movies containing all subreads were used to generate high-fidelity circular consensus sequences (CCS) using pbccs program (https://github.com/PacificBiosciences/ccs) with default settings (minimal number of passages 3, fidelity>98%). CCS reads were demultiplexed and assigned to each sample with the Lima program (https://github.com/pacificbiosciences/barcoding/) using the asymmetric 16 nt barcodes added to the amplicons during PCR amplification (list of barcode combinations per sample in Tables 2-3). Duplicated sequences of the same genomic molecules were removed using the unique molecular identifiers (UMI) of 10 random nucleotides added during SiR genomes retrotransciption. Briefly, UMI tags were extracted from individual reads using UMI_tools (https://github.com/CGATOxford/UMI-tools) and used to generate families of reads from a single original genomic copy. For each family the highest quality read was retained and the others discarded using dedup function of UMI_tools. Deduplicated reads were aligned to the reference using pbmm2 function (https://github.com/PacificBiosciences/pbmm2/) and variants called using the ivar program (https://github.com/andersen-lab/ivar)(Grubaugh et al., 2019) using a minimum base quality of 20. Complete list of the identified mutations and number of reads above q>20 per base per sample can be found in Tables 2 and 3.

#### TEVp-dependency of viral transcription

HEK and HEK-TEVp were seeded in glass bottom wells (μ-Slide 8 Well Glass Bottom, Ibidi) and infected when at ~70% confluence with SiR-nucGFP, SiR-S450X-nucGFP, SiR-G453X-nucGFP or ΔG-Rabies-nucGFP. Live infected cells were imaged 48 hrs post infection in an inverted confocal microscope (SP8 Leica) using a 10x air objective with identical settings for all conditions to evaluate GFP expression levels.

#### Immunohistochemistry

Mice were perfused with ice cold phosphate buffered saline (PBS) followed by 4% paraformaldehyde (PFA) in PBS. Brains were incubated in PFA overnight at 4°C, rinsed twice with PBS followed by dehydration in 30% sucrose in PBS at 4°C for 2 days. Then, brains were frozen in O.C.T. compound (VWR) and sliced at 35 μm on cryostat (Leica, Germany). Free-floating sections were rinsed in PBS and then incubated in blocking solution (1% bovine serum albumin and 0.3% Triton X-100 in PBS) containing primary antibodies for 24 hrs at 4°C. Sections were washed with PBS three times and incubated for 24 hrs at 4°C in blocking solution with secondary antibodies. Immuno-labelled sections were washed three times with PBS and mounted on glass slides. Antibodies used in this study were rabbit anti-RFP (Rockland, 600-401-379, 1:2000) and donkey anti-rabbit Cy3 (Jackson ImmunoResearch, 711-165-152, 1:1000).

#### Viral injections

All procedures using live animals were approved by the Home Office and the LMB Biosafety committee. For all experiments adult mice >8 weeks were used. Mice were anesthetized with 3% isofluorane in 2L/min of oxygen for the initial induction and then maintained with a flow of 1-2% isofluorane in 2 L/min of oxygen. Anesthetized animals were placed into a stereotaxic apparatus (David Kopf Instruments) and Rimadyl (2 mg/kg body weight) was administered subcutaneously (s.c.) as an anti-inflammatory. A small hole (500 μm diameter) was drilled and viruses were injected using a WPI Nanofil syringe (35 gauge) for injections in the hippocampus or a glass capillary for injections in the cerebral cortex. The syringe was left in the brain for 5 min before being retracted. SiR and Rabies viruses were injected at 3-6×10^8^ infectious units/ml. Up to a maximum volume of 500 nl of virus was injected in the following brain areas: hippocampus (AP: −2.45 mm, ML: 2 mm and DV: 1. 5mm from bregma), cerebral cortex (AP: −2. 5 mm, ML: 2 mm and DV: 0,3mm from brain surface).

#### *In vivo* cytotoxicity analysis

SiR-CRE, SiR-G453X-CRE and ΔG-Rabies-CRE *in vivo* cytotoxicity was assessed by injecting 400 nl of purified viral preparations (at 3-6×10^8^ infectious units/ml) in CA1 area of the hippocampus of *Rosa-LoxP-STOP-LoxP-tdTomato* mice. Animals were perfused at 1 week or 1-2 month p.i. and the brains were sectioned at the cryostat (35 μm). The entire hippocampus was sampled (by acquiring one slice every 4) by imaging infected neurons using a robot assisted Nikon HCA microscope mounting a 10x (0.45NA) air objective and tdTomato positive hippocampal neurons counted using Nikon HCA analysis software. Cell survival was calculated by normalizing the total number of infected neurons to the 1 week time point.

#### *In vivo* 2-photon imaging

*Rosa-LoxP-STOP-LoxP-tdTomato* mice aged 3-4 months were injected with Dexafort^®^ at 2 μg/g, one day prior to surgery. Mice were anesthetized with Isofluorane (induction and maintenance at 3% and 2% in 3 L/min of oxygen, respectively) and injected subcutaneously with Vetergesic^®^ at 0.1mg/kg. A metal head-post was affixed to the skull with Crown & Bridge Metabond^®^. Epivicaine^®^ was splashed on the skull, and a 3 mm craniotomy was performed on the left hemisphere, centred at 2mm lateral of the midline and 2,5 mm posterior of bregma. 500 nl of virus with a titer of 4×10^8^ was then delivered at the centre of the craniotomy, at a depth of 300 μm, and at a rate of 100 nl per minute using a manual hydraulic micromanipulator (Narishige). The craniotomy was finally sealed with a 3 mm round coverslip pressing on the brain, and affixed using Crown & Bridge Metabond^®^. Mice were imaged weekly after surgery, under Isofluorane anaesthesia at 1.5% in 3 L/min of oxygen, with a two-photon microscope (Bergamo II, Thorlabs), equipped with a 16x − 0.8 NA objective (Nikon). Infected cells were excited with a Ti:Sapphire pulsed laser at 1030 nm, with a power of around 20mW (Mai Tai DeepSee, Spectra Physics). Emitted fluorescence was collected through a 607 ± 35 nm filter (Brightline). For each mouse, a Z-stack was recorded, centred at the same anterior-posterior coordinate as the injection, but 1mm closer to the midline in the lateral-medial axis. Imaging planes’ pixel resolution was 2048×2048, and depth was sampled in steps of 1 μm. Z-stacks were 3d aligned across time points using a custom program written in Python, segmented into smaller fields of view, and filtered with a 3D mean filter of radius 2 pixels for x and y, and 5 pixels for z (Fiji). All cells at week 1 were labelled using FIJI, and their presence was manually assessed at later time points for the quantification of the survival rate.

#### Quantification and Statistical Analysis

Mean values are accompanied by SEM. No statistical methods were used to predetermine sample sizes. In the hippocampal survival experiments animals were randomly assigned to each time point. Otherwise, data collection and analysis were not performed blind to the conditions of the experiments. Statistical analysis was performed in Graphpad Prism and/or Matlab. Paired t-test and one-way ANOVA test were used to test for statistical significance when appropriate. Statistical parameters including the exact value of n, precision measures (mean ± SEM) and statistical significance are reported in the text and in the figure legends (see individual sections). The significance threshold was placed at α = 0.05.

## Notes

### Competing Interest Statement

The authors have declared no competing interest.

