## Supplementary material for "Genomic stability of Self-inactivating Rabies": Reference sequence for NGS data

LOCUS BSeq#1 1840 bp DNA linear 18-SEP-2020

DEFINITION Synthetic DNA sequence.

ACCESSION BSeq#1

VERSION

KEYWORDS .

SOURCE synthetic DNA sequence

ORGANISM synthetic DNA sequence

Unclassified.

REFERENCE 1 (bases 1 to 1840)

AUTHORS Ciabatti,E.

TITLE Genomic stability of Self-inactivating Rabies

JOURNAL unpublished

REFERENCE 2 (bases 1 to 1840)

AUTHORS Ciabatti,E.

TITLE Direct Submission

JOURNAL Submitted (18-SEP-2020) Neurobiology, MRC - Laboratory of Molecular

Biology, Francis Crick Avenue, Cambridge, Cambridgeshire CB2 0QH,

United Kingdom

COMMENT Bankit Comment: TAX: Yes, new species/combinations; SEE ATTACHMENT

Bankit Comment: TOTAL # OF SEQS:1.

FEATURES Location/Qualifiers

source 1..1840

/organism="synthetic DNA sequence"

/mol_type="other DNA"

gene 72..>1421

/gene="N"

CDS 72..>1421

/gene="N"

/note="Rabies Nucleoprotein"

/codon_start=1

/product="N"

/translation="MDADKIVFKVNNQVVSLKPEIIVDQYEYKYPAIKDLKKPCITLGKAPDL
NKAYKSVLSGMSAAKLNPDDVCSYLAAAMQFFEGTCPEDWTSYGIVIARKGDKITPGS
LVEIKRTDVEGNWALTGGMELTRDPTVPEHASLVGLLLSLYRLSKISGQNTGNYKTNI
ADRIEQIFETAPFVKIVEHHTLMTTHKMCANWSTIPNFRFLAGTYDMFFSRIEHLYSAI
RVGTVVTAYEDCSGLVSFTGFIKQINLTAREAILYFFHKNFEEEIRRMFEPGQETAVPH
SYFIHFRSLGLSGKSPYSSNAVGHVFNLIHFVGCYMGQVRSLNATVIAACAPHEMSVLG
GYLGEEFFGKGTFERRFFRDEKELQEYEAAELTKTDVALADDGTVNSDDEDYFSGETR
SPEAVYTRIMMNGGRLKRSHIRRYVSVSSNHQARPNSFAEFLNKTYSSDS"

CDS <1431..>1451

/note="Tobacco etch virus (TEV) protease cleavage site"

/codon_start=1

/product="TEV site"

/translation="ENLYFQS"

CDS <1461..1583

/note="PEST domain corresponding to residues 422-461 of

mouse ornithine decarboxylase"

/codon_start=1

/product="PEST domain"

/translation="SHGFPPEVEEQDDGTLPMSCAQESGMDRHPAACASARINV"

gene 1674..>1840

/gene="P"

CDS 1674..>1840

/gene="P"

/codon_start=1

/product="P"

/translation="MSKIFVNPSAIRAGLADLEMAEETVDLINRNIEDNQAHLQGEPIEVDN
LPEDMGRL"

BASE COUNT 553 a 394 c 431 g 462 t

ORIGIN

1 cacgcttaac aaccagatca aagaaaaaac agacattgtc aattgcaaag caaaaatgta

61 acacccctac aatggatgcc gacaagattg tattcaaagt caataatcag gtggtctctt

121 tgaagcctga gattatcgtg gatcaatatg agtacaagta ccctgccatc aaagatttga

181 aaaagccctg tataacccta ggaaaggctc ccgatttaaa taaagcatac aagtcagttt

241 tgtcaggcat gagcgccgcc aaacttaatc ctgacgatgt atgttcctat ttggcagcgg

301 caatgcagtt ttttgagggg acatgtccgg aagactggac cagctatgga attgtgattg

361 cacgaaaagg agataagatc accccaggtt ctctggtgga gataaaacgt actgatgtag

421 aagggaattg ggctctgaca ggaggcatgg aactgacaag agaccccact gtccctgagc

481 atgcgtcctt agtcggtctt ctcttgagtc tgtataggtt gagcaaaata tccgggcaaa

541 acactggtaa ctataagaca aacattgcag acaggataga gcagattttt gagacagccc

601 cttttgttaa aatcgtggaa caccatactc taatgacaac tcacaaaatg tgtgctaatt

661 ggagtactat accaaacttc agatttttgg ccggaaccta tgacatgttt ttctcccgga

721 ttgagcatct atattcagca atcagagtgg gcacagttgt cactgcttat gaagactgtt

781 caggactggt atcatttact gggttcataa aacaaatcaa tctcaccgct agagaggcaa

841 tactatattt cttccacaag aactttgagg aagagataag aagaatgttt gagccagggc

901 aggagacagc tgttcctcac tcttatttca tccacttccg ttcactaggc ttgagtggga

961 aatctcctta ttcatcaaat gctgttggtc acgtgttcaa tctcattcac tttgtaggat

1021 gctatatggg tcaagtcaga tccctaaatg caacggttat tgctgcatgt gctcctcatg

1081 aaatgtctgt tctagggggc tatctgggag aggaattctt cgggaaaggg acatttgaaa

1141 gaagattctt cagagatgag aaagaacttc aagaatacga ggcggctgaa ctgacaaaga

1201 ctgacgtagc actggcagat gatggaactg tcaactctga cgacgaggac tacttttcag

1261 gtgaaaccag aagtccggag gctgtttata ctcgaatcat gatgaatgga ggtcgactaa

1321 agagatctca catacggaga tatgtctcag tcagttccaa tcatcaagcc cgtccaaact

1381 cattcgccga gtttctaaac aagacatatt cgagtgactc aggttccgga gagaacctct

1441 acttccaatc gggatccggt agccatggct tcccgccgga ggtggaggag caggatgatg

1501 gcacgctgcc catgtcttgt gcccaggaga gcgggatgga ccgtcaccct gcagcctgtg

1561 cttctgctag gatcaatgtg taagaagttg aataacaaaa tgccggaaat ctacggattg

1621 tgtatatcca tcatgaaaaa aactaacacc cctcctttcg aaccatccca aacatgagca

1681 agatctttgt caatcctagt gctattagag ccggtctggc cgatcttgag atggctgaag

1741 aaactgttga tctgatcaat agaaatatcg aagacaatca ggctcatctc caaggggaac

1801 ccatagaggt ggacaatctc cctgaggata tggggcgact

//
