## Supplementary Figures for "Genomic stability of Self-inactivating Rabies"

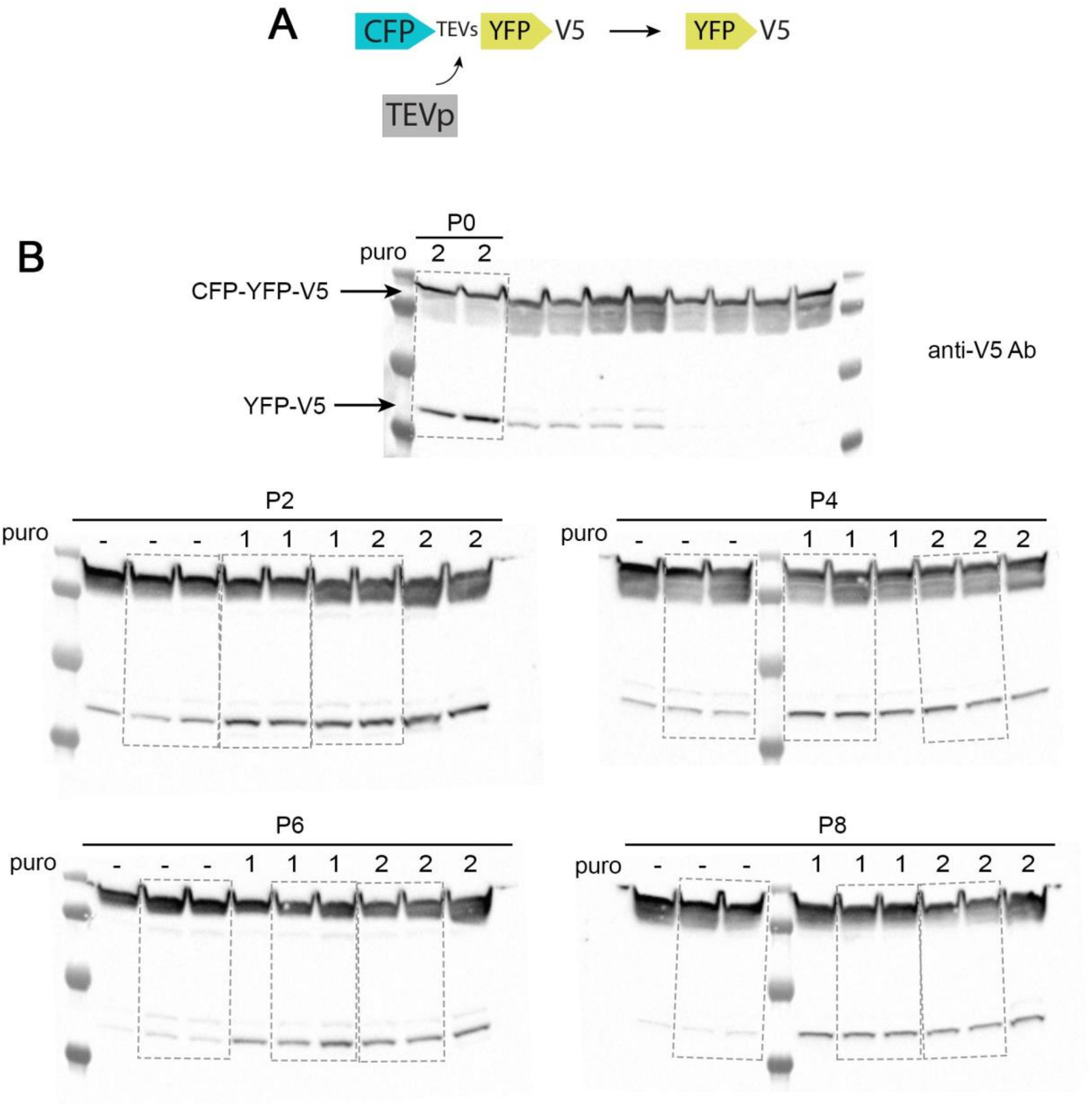


**Fig S1 Western blots to test TEVp in packaging cells over time**

A) Scheme of the TEVp activity reporter. B) Original western blots stained with an anti-V5 antibody with the representative lanes used to generate figure 2B.

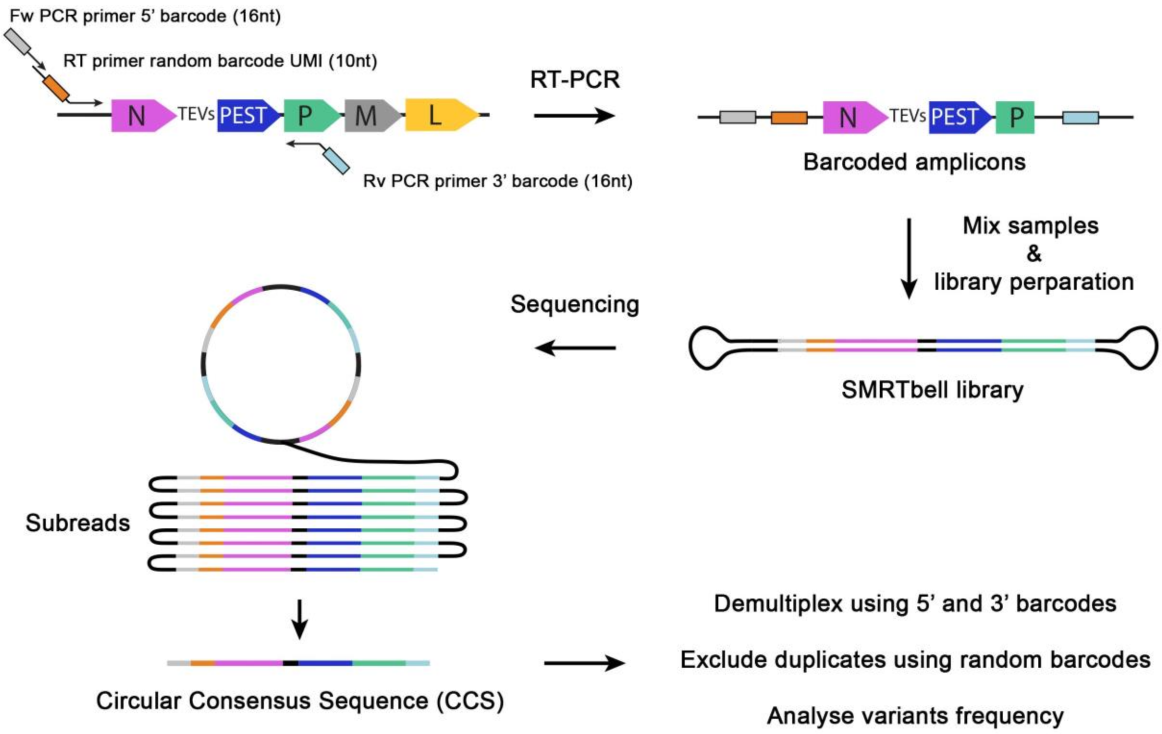


**Fig S2 SMRT sequencing of SiR genomic libraries**

Scheme of the strategy to sequence SiR preparations using SMRT NGS technology from Pacbio. Amplicons of the entire coding sequence of N-TEVs-PEST gene are generated by RT-PCR. Unique Molecular Identifier (UMI) of 10 nucleotides is added during retrotranscription to each genomic molecule and sample specific barcodes of 16 nucleotides are added at the two ends during subsequent PCR. SMRT bell libraries are generated by ligating the provided adapters to generate circular DNA molecules that are sequenced continuously for multiple passages. Subreads are used to generate high-fidelity consensus sequences that are demultiplexed using the 16 nt barcodes, deduplicated using the UMIs and aligned to the reference for variant calling.

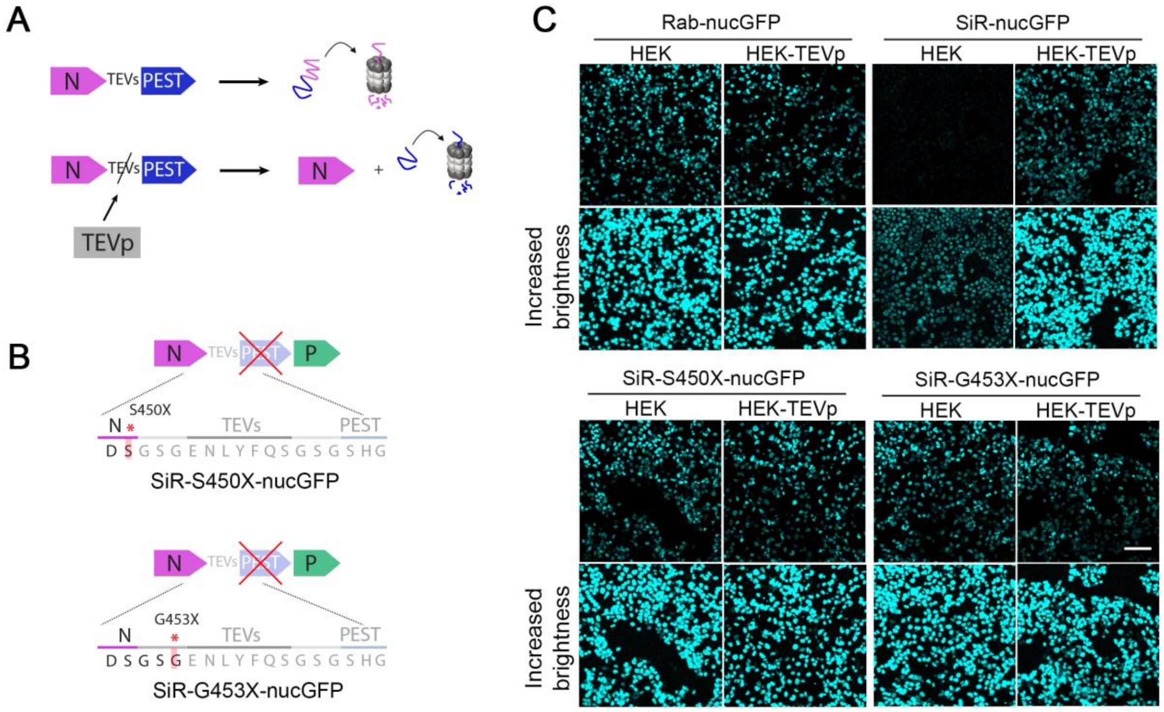


**Fig S3 SiR revertants lose functional TEVs and PEST domain**

A) The conditional destabilization of N can be prevented by TEVp expression in the infected cells leading to cleavage of the TEVs-containing linker. B) Engineered revertant SiR viruses containing the reporter PEST-inactivating substitutions in their cDNA. C) Confocal images of HEK and HEK-TEVp at 48 hrs p.i. All images were acquired with same settings. Bottom panels have been equally adjusted in brightness in all conditions. Scale bar 100 μm.
