## Supplementary material for "Genomic stability of Self-inactivating Rabies": Table 1

### Sanger sequencing results of SiRs rescued from cDNA

| Batch A | | | | | |
|---|---|---|---|---|---|
| | Clones | Sequence | Position | Mutation | Effect on CDS |
| Upstream N | 1/50 | GAT > GAC | -54 | Substitution | - |
| | 1/50 | AAA > AAG | -18 | Substitution | - |
| N gene | 1/50 | GCC > GCT | +186 | Substitution | Synonymous A62 |
| | 1/50 | TTT > TTTT | +243 | Insertion | Frameshift |
| | 1/50 | AAG > A-G | +485 | Deletion | Frameshift |
| | 1/50 | ATG > CTG | +562 | Substitution | Missense M188L |
| | 1/50 | GTG > G-- | +677/8 | Deletion | Frameshift |
| | 1/50 | ACG > ACCG | +983 | Insertion | Frameshift |
| | 1/50 | GAA > AAA | +1093 | Substitution | E365K |
| | 1/50 | TCA > CCA | +1276 | Substitution | S426P |
| TEVs-PEST | - | - | - | - | - |
| Intergenic N/P | 4/50 | AAA > AAAA | +1571 | Insertion | - |
| | 1/50 | CCC > CCA | +1581 | Substitution | - |
| P | - | - | - | - | - |

| Batch B | | | | | |
|---|---|---|---|---|---|
| | Clones | Sequence | Position | Mutation | Effect on CDS |
| Upstream N | 1/50 | AAC > A-C | -63 | Deletion | - |
| | 1/50 | CAA > CA- | -60 | Deletion | |
| | 1/50 | CTA > CTG | -3 | Substitution | - |
| N gene | 1/50 | TTT > TTTT | +243 | Insertion | Frameshift |
| | 1/50 | GAC > GAA | +501 | Substitution | D167E |
| | 1/50 | AAT > AAC | +588 | Substitution | Synonymous N196 |
| | 1/50 | GCT > GCC | +1002 | Substitution | Synonymous A334 |
| | 1/50 | AAA > AAAA | +1056 | Insertion | Frameshift |
| TEVs-PEST | 1/50 | TCC > TGC | +1385 | Substitution | Missense S462C in GSG linker after TEVs |
| Intergenic N/P | 1/50 | TAT > TAA | +1554 | Substitution | - |
| | 2/50 | AAA > AAAA | +1571 | Insertion | - |
| P | 1/50 | GAA > GAG | +1671 | Substitution | Synonymous E23 |
| | | CTG > CCG | +1775 | Substitution | Missense L58P |
| | | GGA > TGA | +2014 | Deletion | Nonsense G138>STOP |

| Batch C | | | | | |
|---|---|---|---|---|---|
| | Clones | Sequence | Position | Mutation | Effect on CDS |
| Upstream N | 2/50 | AAA > AAAA | -43 | Insertion | - |
| N gene | 1/50 | TGT > TTT | +212 | Substitution | Missense C71F |
| | 1/50 | AGA > AGG | +1074 | Substitution | Synonymous R358 |
| | 1/50 | GGT > GAT | +1190 | Substitution | Missense G397D |
| TEVs-PEST | - | - | - | - | - |
| Intergenic N/P | 1/50 | AAA > AAG | +1569 | Substitution | - |

| | | | | | |
|---|---|---|---|---|---|
| | 3/50 | AAA > AAAA | +1571 | Insertion | - |
| | 1/50 | AAA > AA- | +1571 | Deletion | |
| P | 1/50 | CAA > AAA | +1720 | Substitution | Missense Q40K |

| Batch D | | | | | |
|---|---|---|---|---|---|
| | Clones | Sequence | Position | Mutation | Effect on CDS |
| Upstream N | - | - | - | - | - |
| N gene | 1/50 | AAG > AGG | +113 | Substitution | Missense K38R |
| | 1/50 | AAA > CAA | +295 | Substitution | Missense K99Q |
| | 1/50 | CAT > AAT | +655 | Substitution | Missense H219N |
| | 1/50 | TCA > TCC | +873 | Substitution | Synonymous S291 |
| | 1/50 | ACC > AAC | +1196 | Substitution | Missense T399N |
| TEVs-PEST | - | - | - | - | - |
| Intergenic N/P | 3/50 | AAA > AAAA | +1571 | Insertion | - |
| | 1/50 | ATC > ATT | +1596 | Substitution | - |
| P | 1/50 | AAA > AAAA | +1671 | Insertion | Frameshift |
| | 1/50 | CGT > CTA | +1878 | Substitution | Synonymous L92 |
| | 1/50 | AGA > AGT | +1941 | Substitution | Missense R113S |
| | 1/50 | GGA > GGG | +2016 | Substitution | Synonymous G138 |
| | 1/50 | ACT > ACA | +2046 | Substitution | Synonymous T148 |

| Batch E | | | | | |
|---|---|---|---|---|---|
| | Clones | Sequence | Position | Mutation | Effect on CDS |
| Upstream N | 1/50 | CCA > CC- | -57 | Deletion | - |
| N gene | 1/50 | CCT >CAT | +200 | Substitution | Missense P67H |
| | 1/50 | TTT > TTTT | +243 | Insertion | Frameshift |
| | 1/50 | GGA > GAA | +371 | Substitution | Missense G124E |
| | 1/50 | ACA > ACG | +387 | Substitution | Synonymous T129 |
| | 2/50 | GAC > GAT | +393 | Substitution | Synonymous D131 |
| | 1/50 | CAC > C-- | +551/2 | Deletion | Frameshift |
| | 1/50 | ACT > AAT | +557 | Substitution | T186N |
| | 1/50 | TTT > TTTT | +779 | Insertion | Frameshift |
| TEVs-PEST | - | - | - | - | - |
| Intergenic N/P | 1/50 | CAT > CAC | +1560 | Substitution | - |
| | 1/50 | AAA > AAC | +1570 | Substitution | |
| | 4/50 | AAA > AAAA | +1571 | Insertion | |
| | 1/50 | ATC > ATT | +1596 | Substitution | - |
| P | 1/50 | GAA > GGA | +1667 | Substitution | Missense E22G |

| Batch F | | | | | |
|---|---|---|---|---|---|
| | Clones | Sequence | Position | Mutation | Effect on CDS |
| Upstream N | 1/50 | ACC > AC- | -58 | Deletion | - |
| | 1/50 | CAG > CA- | -56 | Deletion | - |
| | 1/50 | TCA > TCG | -52 | Substitution | - |
| | 1/50 | AAA > AAAA | -43 | Insertion | - |
| | 1/50 | AAG > AA- | -22 | Deletion | - |
| N gene | 1/50 | TTT >TTTTT | +243/4 | Insertion | Frameshift |
| | 1/50 | TTG > TCG | +434 | Substitution | Missense L145S |

| | | | | | |
|---|---|---|---|---|---|
| | 1/50 | TTT > TT- | +534 | Deletion | Frameshift |
| | 1/50 | GCA > GTA | +767 | Substitution | Missense A256V |
| | 1/50 | ACA > ATA | +836 | Substitution | Missense T279I |
| | 1/50 | AAA > AAAA | +908 | Insertion | Frameshift |
| | 1/50 | 321 bp | +1041 - 1362 | Deletion | Deletion of C-terminal of N in frame with PEST domain |
| | 1/50 | GGA > GGG | +1038 | Substitution | Synonymous G346 |
| TEVs-PEST | - | - | - | - | - |
| Intergenic N/P | 4/50 | AAA > AAAA | +1571 | Insertion | - |
| P | 1/50 | CCT > CCC | +1626 | Substitution | Synonymous P8 |
| | 1/50 | GAA > GGA | +1727 | Substitution | Missense E42G |
| | 1/50 | TTT > TTC | +1845 | Substitution | Synonymous F81 |

| Batch G | | | | | |
|---|---|---|---|---|---|
| | Clones | Sequence | Position | Mutation | Effect on CDS |
| Upstream N | 1/50 | CCA > CC- | -57 | Deletion | - |
| | 1/50 | AAA > AA- | -16 | Deletion | - |
| N gene | 1/50 | GCA > GTA | +290 | Substitution | Missense A97V |
| | 1/50 | CAT > GAT | +409 | Substitution | Missense H137D |
| | 1/50 | TTT > TT- | +534 | Deletion | Frameshift |
| | 1/50 | TAT > TGT | +1271 | Substitution | Missense Y424C |
| | 1/50 | GCC > GTC | +1316 | Substitution | Missense A439V |
| TEVs-PEST | - | - | - | - | - |
| Intergenic N/P | 4/50 | AAA > AAAA | +1571 | Insertion | - |
| P | 1/50 | AAA > CAA | +1786 | Substitution | Missense K62Q |
| | 1/50 | GAA > GGA | +1823 | Substitution | Missense E74G |
| | 1/50 | CGA > CAA | +1834 | Substitution | Missense R78Q |

| Batch H | | | | | |
|---|---|---|---|---|---|
| | Clones | Sequence | Position | Mutation | Effect on CDS |
| Upstream N | 1/50 | AAA > AAAA | -43 | Insertion | - |
| | 1/50 | AAC > AA- | -42 | Deletion | |
| N gene | 1/50 | TTA > CTA | +145 | Substitution | Synonymous L49 |
| | 1/50 | ATG > ATA | +234 | Substitution | Missense M78I |
| | 1/50 | TTT > TTTT | +243 | Insertion | Frameshift |
| | 1/50 | AAA > CAA | +295 | Substitution | Missense K99Q |
| | 1/50 | GAT > AAT | +301 | Substitution | Missense D101N |
| | 1/50 | GGA > AGA | +622 | Substitution | Missense G208R |
| | 1/50 | GCT > TCT | +838 | Substitution | Missense A280S |
| | 1/50 | GGC > G-C | +1028 | Deletion | Frameshift |
| | 1/50 | GAC > AAC | +1132 | Substitution | Missense D378N |
| TEVs-PEST | 1/50 | CTG > CTA | +1437 | Substitution | Synonymous L16 in PEST domain |
| Intergenic N/P | 3/50 | AAA > AAAA | +1571 | Insertion | - |
| | 1/50 | AAC > AAA | +1592 | Substitution | - |
| P | 1/50 | AAA > AAAA | +1788 | Insertion | Frameshift |

**Table 1**

List of detected mutations in SiR viruses rescued from cDNA divided by batch (50 individual clones per batch). The position of the mutations is calculated referring to +1 as the first base of the nucleoprotein N coding sequence.
