## Supplementary material for "Genomic stability of Self-inactivating Rabies": Table 2

### NGS sequencing results of SiRs amplified for multiple passages *in vitro*

| SIR-A-P0 bc1—bc2 | | | | | | |
|---|---|---|---|---|---|---|
| | Position | Variant | N (q>20) | Freq % | Mutation | Effect on CDS |
| Upstream N | -49 | +A | 302/6608 | 4.5% | Insertion | - |
| N gene | +237 | +T | 266/6598 | 4.0% | Insertion | Frameshift |
| | +636 | +T | 190/6595 | 2.9% | Insertion | Frameshift |
| TEVs-PEST | - | - | - | - | - | - |
| Intergenic | +1564 | +A | 732/6556 | 11.1% | Insertion | - |
| P | - | - | - | - | - | - |

| SIR-B-P0 bc1—bc3 | | | | | | |
|---|---|---|---|---|---|---|
| | Position | Variant | N (q>20) | Freq % | Mutation | Effect on CDS |
| Upstream N | -49 | +A | 276/6045 | 4.6% | Insertion | - |
| N gene | +237 | +T | 274/6037 | 4.5% | Insertion | Frameshift |
| | +636 | +T | 180/6036 | 3.0% | Insertion | Frameshift |
| TEVs-PEST | +1359 | A>T | 246/5879 | 4.2% | Substitution | Silent G453 |
| Intergenic | +1564 | +A | 729/6556 | 12.1% | Insertion | - |
| P | - | - | - | - | - | - |

| SIR-C-P0 bc1—bc4 | | | | | | |
|---|---|---|---|---|---|---|
| | Position | Variant | N (q>20) | Freq % | Mutation | Effect on CDS |
| Upstream N | -49 | +A | 256/5137 | 5.0% | Insertion | - |
| N gene | +237 | +T | 227/5137 | 4.4% | Insertion | Frameshift |
| | +636 | +T | 167/5138 | 3.3% | Insertion | Frameshift |
| TEVs-PEST | - | - | - | - | - | - |
| Intergenic | +1564 | +A | 598/5140 | 11.6% | Insertion | - |
| P | - | - | - | - | - | - |

| SIR-D-P0 bc1—bc5 | | | | | | |
|---|---|---|---|---|---|---|
| | Position | Variant | N (q>20) | Freq % | Mutation | Effect on CDS |
| Upstream N | -49 | +A | 249/5419 | 4.6% | Insertion | - |
| N gene | +237 | +T | 229/5419 | 4.2% | Insertion | Frameshift |
| | +636 | +T | 125/5422 | 2.3% | Insertion | Frameshift |
| TEVs-PEST | - | - | - | - | - | - |
| Intergenic | +1564 | +A | 612/5420 | 11.3% | Insertion | - |
| P | - | - | - | - | - | - |

| SIR-A-HighTEVp-P2 bc2—bc4 | | | | | | |
|---|---|---|---|---|---|---|
| | Position | Variant | N (q>20) | Freq % | Mutation | Effect on CDS |
| Upstream N | -49 | +A | 245/5934 | 4.1% | Insertion | - |
| N gene | +237 | +T | 297/5933 | 5.0% | Insertion | Frameshift |
| | +636 | +T | 157/5938 | 2.6% | Insertion | Frameshift |
| TEVs-PEST | - | - | - | - | - | - |
| Intergenic | +1564 | +A | 634/5935 | 10.7% | Insertion | - |

| | | | | | | |
|---|---|---|---|---|---|---|
| P | - | - | - | - | - | - |

| SIR-B-HighTEVp-P2 bc2—bc5 | | | | | | |
|---|---|---|---|---|---|---|
| | Position | Variant | N (q>20) | Freq % | Mutation | Effect on CDS |
| Upstream N | -49 | +A | 281/5750 | 4.9% | Insertion | - |
| N gene | +237 | +T | 272/5752 | 4.7% | Insertion | Frameshift |
| | +636 | +T | 170/5752 | 3.0% | Insertion | Frameshift |
| TEVs-PEST | - | - | - | - | - | - |
| Intergenic | +1564 | +A | 625/5749 | 10.9% | Insertion | - |
| P | - | - | - | - | - | - |

| SIR-C-HighTEVp-P2 bc2—bc6 | | | | | | |
|---|---|---|---|---|---|---|
| | Position | Variant | N (q>20) | Freq % | Mutation | Effect on CDS |
| Upstream N | -49 | +A | 236/4773 | 4.9% | Insertion | - |
| N gene | +237 | +T | 241/4772 | 5.1% | Insertion | Frameshift |
| | +636 | +T | 137/4774 | 2.9% | Insertion | Frameshift |
| TEVs-PEST | - | - | - | - | - | - |
| Intergenic | +1564 | +A | 489/4776 | 10.2% | Insertion | - |
| P | - | - | - | - | - | - |

| SIR-D-HighTEVp-P2 bc2—bc6 | | | | | | |
|---|---|---|---|---|---|---|
| | Position | Variant | N (q>20) | Freq % | Mutation | Effect on CDS |
| Upstream N | -49 | +A | 260/5591 | 4.7% | Insertion | - |
| N gene | +237 | +T | 238/5595 | 4.3% | Insertion | Frameshift |
| | +636 | +T | 150/5597 | 2.7% | Insertion | Frameshift |
| TEVs-PEST | - | - | - | - | - | - |
| Intergenic | +1564 | +A | 550/5594 | 9.8% | Insertion | - |
| P | - | - | - | - | - | - |

| SIR-A-LowTEVp-P2 bc1—bc6 | | | | | | |
|---|---|---|---|---|---|---|
| | Position | Variant | N (q>20) | Freq % | Mutation | Effect on CDS |
| Upstream N | -49 | +A | 197/3891 | 5.1% | Insertion | - |
| N gene | +237 | +T | 194/3891 | 5.0% | Insertion | Frameshift |
| | +636 | +T | 116/3892 | 3.0% | Insertion | Frameshift |
| TEVs-PEST | - | - | - | - | - | - |
| Intergenic | +1564 | +A | 447/3891 | 11.5% | Insertion | - |
| P | - | - | - | - | - | - |

| SIR-B-LowTEVp-P2 bc1—bc7 | | | | | | |
|---|---|---|---|---|---|---|
| | Position | Variant | N (q>20) | Freq % | Mutation | Effect on CDS |
| Upstream N | -49 | +A | 244/5050 | 4.8% | Insertion | - |
| N gene | +237 | +T | 227/5055 | 4.5% | Insertion | Frameshift |
| | +636 | +T | 162/5055 | 3.2% | Insertion | Frameshift |
| TEVs-PEST | - | - | - | - | - | - |
| Intergenic | +1564 | +A | 503/5055 | 10.0% | Insertion | - |
| P | - | - | - | - | - | - |

| SIR-C-LowTEVp-P2 bc1—bc8 | | | | | | |
|---|---|---|---|---|---|---|
| | Position | Variant | N (q>20) | Freq % | Mutation | Effect on CDS |
| Upstream N | -49 | +A | 266/5050 | 5.3% | Insertion | - |
| N gene | +237 | +T | 248/5050 | 4.9% | Insertion | Frameshift |
| | +636 | +T | 146/5056 | 2.9% | Insertion | Frameshift |
| TEVs-PEST | - | - | - | - | - | - |
| Intergenic | +1564 | +A | 547/5054 | 10.8% | Insertion | - |
| P | - | - | - | - | - | - |

| SIR-D-LowTEVp-P2 bc1—bc9 | | | | | | |
|---|---|---|---|---|---|---|
| | Position | Variant | N (q>20) | Freq % | Mutation | Effect on CDS |
| Upstream N | -49 | +A | 200/5295 | 3.8% | Insertion | - |
| N gene | +237 | +T | 204/5295 | 3.9% | Insertion | Frameshift |
| | +636 | +T | 141/5297 | 2.7% | Insertion | Frameshift |
| TEVs-PEST | - | - | - | - | - | - |
| Intergenic | +1564 | +A | 456/5297 | 8.6% | Insertion | - |
| P | - | - | - | - | - | - |

| SIR-A-HighTEVp-P4 bc2—bc8 | | | | | | |
|---|---|---|---|---|---|---|
| | Position | Variant | N (q>20) | Freq % | Mutation | Effect on CDS |
| Upstream N | -49 | +A | 225/5803 | 3.9% | Insertion | - |
| N gene | +108 | +A | 154/5805 | 2.7% | Insertion | Frameshift |
| | +237 | +T | 276/5806 | 4.8% | Insertion | Frameshift |
| | +636 | +T | 158/5807 | 2.7% | Insertion | Frameshift |
| TEVs-PEST | +1357 | G>T | 134/5745 | 2.3% | Substitution | Missense G453X |
| Intergenic | +1564 | +A | 536/5803 | 9.2% | Insertion | - |
| P | - | - | - | - | - | - |

| SIR-B-HighTEVp-P4 bc2—bc10 | | | | | | |
|---|---|---|---|---|---|---|
| | Position | Variant | N (q>20) | Freq % | Mutation | Effect on CDS |
| Upstream N | -49 | +A | 270/5572 | 4.8% | Insertion | - |
| N gene | +237 | +T | 223/5572 | 4.0% | Insertion | Frameshift |
| | +636 | +T | 155/5571 | 2.8% | Insertion | Frameshift |
| TEVs-PEST | - | - | - | - | - | - |
| Intergenic | +1564 | +A | 590/5576 | 10.6% | Insertion | - |
| P | - | - | - | - | - | - |

| SIR-C-HighTEVp-P4 bc2—bc11 | | | | | | |
|---|---|---|---|---|---|---|
| | Position | Variant | N (q>20) | Freq % | Mutation | Effect on CDS |
| Upstream N | -49 | +A | 233/5581 | 4.2% | Insertion | - |
| | -21 | -N | 114/5581 | 2.0% | Deletion | - |
| | -19 | A>G | 272/5499 | 4.9% | Substitution | - |
| N gene | +237 | +T | 252/5582 | 4.5% | Insertion | Frameshift |
| | +636 | +T | 149/5581 | 2.7% | Insertion | Frameshift |

| | | | | | | |
|---|---|---|---|---|---|---|
| TEVs-PEST | +1357 | G>T | 248/5528 | 4.5% | Substitution | Missense G453X |
| Intergenic | +1564 | +A | 573/5579 | 10.3% | Insertion | - |
| P | - | - | - | - | - | - |

| SIR-D-HighTEVp-P4 bc2—bc12 | | | | | | |
|---|---|---|---|---|---|---|
| | Position | Variant | N (q>20) | Freq % | Mutation | Effect on CDS |
| Upstream N | -49 | +A | 200/6116 | 3.3% | Insertion | - |
| N gene | +237 | +T | 219/6117 | 3.6% | Insertion | Frameshift |
| | +636 | +T | 160/6119 | 2.6% | Insertion | Frameshift |
| TEVs-PEST | - | - | - | - | - | - |
| Intergenic | +1564 | +A | 456/6120 | 7.5% | Insertion | - |
| P | - | - | - | - | - | - |

| SIR-A-LowTEVp-P4 bc1—bc10 | | | | | | |
|---|---|---|---|---|---|---|
| | Position | Variant | N (q>20) | Freq % | Mutation | Effect on CDS |
| Upstream N | -49 | +A | 239/4681 | 5.1% | Insertion | - |
| N gene | +108 | +A | 114/4682 | 2.4% | Insertion | Frameshift |
| | +237 | +T | 242/4683 | 5.2% | Insertion | Frameshift |
| | +636 | +T | 131/4684 | 2.8% | Insertion | Frameshift |
| | +1053 | +A | 97/4683 | 2.1% | Insertion | Frameshift |
| TEVs-PEST | +1357 | G>T | 170/4650 | 3.7% | Substitution | Missense G453X |
| Intergenic | +1564 | +A | 570/4683 | 12.2% | Insertion | - |
| P | - | - | - | - | - | - |

| SIR-B-LowTEVp-P4 bc1—bc11 | | | | | | |
|---|---|---|---|---|---|---|
| | Position | Variant | N (q>20) | Freq % | Mutation | Effect on CDS |
| Upstream N | -49 | +A | 255/4757 | 5.4% | Insertion | - |
| N gene | +237 | +T | 245/4758 | 5.1% | Insertion | Frameshift |
| | +636 | +T | 141/4758 | 3.0% | Insertion | Frameshift |
| TEVs-PEST | - | - | - | - | - | - |
| Intergenic | +1564 | +A | 551/4757 | 11.6% | Insertion | - |
| P | - | - | - | - | - | - |

| SIR-C-LowTEVp-P4 bc1—bc12 | | | | | | |
|---|---|---|---|---|---|---|
| | Position | Variant | N (q>20) | Freq % | Mutation | Effect on CDS |
| Upstream N | -49 | +A | 268/5461 | 4.9% | Insertion | - |
| | -19 | A>G | 160/5403 | 3.0% | Substitution | - |
| N gene | +237 | +T | 231/5463 | 4.2% | Insertion | Frameshift |
| | +636 | +T | 156/5466 | 2.9% | Insertion | Frameshift |
| TEVs-PEST | +1357 | G>T | 705/5286 | 13.3% | Substitution | Missense G453X |
| Intergenic | +1564 | +A | 538/5464 | 9.8% | Insertion | - |
| P | - | - | - | - | - | - |

| SIR-D-LowTEVp-P4 bc2—bc3 | | | | | | |
|---|---|---|---|---|---|---|
| | Position | Variant | N (q>20) | Freq % | Mutation | Effect on CDS |

| | | | | | | |
|---|---|---|---|---|---|---|
| Upstream N | -49 | +A | 266/5841 | 4.6% | Insertion | - |
| N gene | +237 | +T | 246/5838 | 4.2% | Insertion | Frameshift |
| | +574 | -N | 140/5834 | 2.4% | Deletion | Frameshift |
| | +636 | +T | 156/5833 | 2.7% | Insertion | Frameshift |
| TEVs-PEST | +1357 | G>T | 200/5737 | 3.5% | Substitution | Missense G453X |
| Intergenic | +1564 | +A | 529/5818 | 9.1% | Insertion | - |
| P | - | - | - | - | - | - |

| SIR-A-HighTEVp-P6 bc5—bc6 | | | | | | |
|---|---|---|---|---|---|---|
| | Position | Variant | N (q>20) | Freq % | Mutation | Effect on CDS |
| Upstream N | -49 | +A | 604/6567 | 9.2% | Insertion | - |
| | -19 | A>G | 555/6349 | 8.7% | Substitution | - |
| N gene | +108 | +A | 227/6565 | 3.5% | Insertion | Frameshift |
| | +166 | +T | 157/6565 | 2.4% | Insertion | Frameshift |
| | +237 | +T | 543/6565 | 8.3% | Insertion | Frameshift |
| | +245 | +G | 132/6565 | 2.0% | Insertion | Frameshift |
| | +466 | +A | 175/6566 | 2.7% | Insertion | Frameshift |
| | +636 | +T | 337/6569 | 5.1% | Insertion | Frameshift |
| TEVs-PEST | +1357 | G>T | 767/6317 | 12.1% | Substitution | Missense G453X |
| Intergenic | +1564 | +A | 1032/6583 | 15.7% | Insertion | - |
| P | +1669 | +A | 155/6584 | 2.4% | Insertion | Frameshift |

| SIR-B-HighTEVp-P6 bc5—bc7 | | | | | | |
|---|---|---|---|---|---|---|
| | Position | Variant | N (q>20) | Freq % | Mutation | Effect on CDS |
| Upstream N | -49 | +A | 624/6752 | 9.2% | Insertion | - |
| | -21 | -N | 202/6754 | 3.0% | Deletion | - |
| | -20 | +G | 243/6754 | 3.6% | Insertion | - |
| | -19 | A>G | 1180/6296 | 18.7% | Substitution | - |
| N gene | +108 | +A | 216/6752 | 3.2% | Insertion | Frameshift |
| | +166 | +T | 185/6751 | 2.7% | Insertion | Frameshift |
| | +237 | +T | 559/6751 | 8.3% | Insertion | Frameshift |
| | +245 | +G | 138/6751 | 2.0% | Insertion | Frameshift |
| | +466 | +A | 197/6753 | 2.9% | Insertion | Frameshift |
| | +612 | +T | 147/6753 | 2.2% | Insertion | Frameshift |
| | +636 | +T | 330/6753 | 4.9% | Insertion | Frameshift |
| TEVs-PEST | - | - | - | - | - | - |
| Intergenic | +1564 | +A | 965/6766 | 14.3% | Insertion | - |
| P | +1669 | +A | 187/6769 | 2.8% | Insertion | Frameshift |

| SIR-C-HighTEVp-P6 bc5—bc8 | | | | | | |
|---|---|---|---|---|---|---|
| | Position | Variant | N (q>20) | Freq % | Mutation | Effect on CDS |
| Upstream N | -49 | +A | 578/6166 | 9.4% | Insertion | - |
| | -21 | -N | 205/6166 | 3.3% | Deletion | - |
| | -20 | +G | 298/6166 | 4.8% | Insertion | - |
| | -19 | A>G | 3305/5625 | 58.8% | Substitution | - |

| | | | | | | |
|---|---|---|---|---|---|---|
| N gene | +108 | +A | 179/6166 | 2.9% | Insertion | Frameshift |
| | +166 | +T | 171/6165 | 2.8% | Insertion | Frameshift |
| | +237 | +T | 514/6164 | 8.3% | Insertion | Frameshift |
| | +466 | +A | 158/6166 | 2.6% | Insertion | Frameshift |
| | +636 | +T | 318/6170 | 5.2% | Insertion | Frameshift |
| TEVs-PEST | +1357 | G>T | 436/5995 | 7.3% | Substitution | Missense G453X |
| Intergenic | +1564 | +A | 1019/6184 | 16.5% | Insertion | - |
| P | +1669 | +A | 165/6185 | 2.7% | Insertion | Frameshift |

| SIR-D-HighTEVp-P6 bc5—bc9 | | | | | | |
|---|---|---|---|---|---|---|
| | Position | Variant | N (q>20) | Freq % | Mutation | Effect on CDS |
| Upstream N | -49 | +A | 562/6355 | 8.8% | Insertion | - |
| | -21 | -N | 228/6356 | 3.6% | Deletion | - |
| | -20 | +G | 314/6356 | 4.9% | Insertion | - |
| | -19 | A>G | 2816/5789 | 48.6% | Substitution | - |
| | -9 | A>T | 139/6104 | 2.3% | Substitution | - |
| | -6 | C>T | 176/6275 | 2.8% | Substitution | - |
| | -5 | C>A | 121/5995 | 2.0% | Substitution | - |
| N gene | +108 | +A | 175/6357 | 2.8% | Insertion | Frameshift |
| | +237 | +T | 474/6358 | 7.5% | Insertion | Frameshift |
| | +245 | +G | 131/6358 | 2.1% | Insertion | Frameshift |
| | +466 | +A | 167/6359 | 2.6% | Insertion | Frameshift |
| | +636 | +T | 316/6360 | 5.0% | Insertion | Frameshift |
| TEVs-PEST | - | - | - | - | - | - |
| Intergenic | +1564 | +A | 947/6365 | 14.9% | Insertion | - |
| P | +1669 | +A | 139/6365 | 2.2% | Insertion | Frameshift |

| SIR-A-LowTEVp-P6 bc4—bc5 | | | | | | |
|---|---|---|---|---|---|---|
| | Position | Variant | N (q>20) | Freq % | Mutation | Effect on CDS |
| Upstream N | -49 | +A | 588/6703 | 8.8% | Insertion | - |
| | -19 | A>G | 369/6525 | 5.7% | Substitution | - |
| N gene | +108 | +A | 259/6704 | 3.9% | Insertion | Frameshift |
| | +166 | +T | 173/6704 | 2.6% | Insertion | Frameshift |
| | +237 | +T | 584/6703 | 8.7% | Insertion | Frameshift |
| | +246 | +G | 145/6703 | 2.2% | Insertion | Frameshift |
| | +466 | +A | 196/6704 | 2.9% | Insertion | Frameshift |
| | +636 | +T | 366/6705 | 5.5% | Insertion | Frameshift |
| TEVs-PEST | +1357 | G>T | 681/6468 | 10.5% | Substitution | Missense G453X |
| Intergenic | +1564 | +A | 1035/6711 | 15.4% | Insertion | - |
| P | +1669 | +A | 161/6711 | 2.4% | Insertion | Frameshift |

| SIR-B-LowTEVp-P6 bc4—bc6 | | | | | | |
|---|---|---|---|---|---|---|
| | Position | Variant | N (q>20) | Freq % | Mutation | Effect on CDS |
| Upstream N | -49 | +A | 550/6112 | 9.0% | Insertion | - |
| | -19 | A>G | 317/5985 | 5.3% | Substitution | - |

| | | | | | | |
|---|---|---|---|---|---|---|
| N gene | +108 | +A | 186/6117 | 3.0% | Insertion | Frameshift |
| | +166 | +T | 131/6117 | 2.1% | Insertion | Frameshift |
| | +237 | +T | 486/6116 | 7.9% | Insertion | Frameshift |
| | +466 | +A | 148/6118 | 2.4% | Insertion | Frameshift |
| | +612 | +T | 125/6120 | 2.0% | Insertion | Frameshift |
| | +636 | +T | 303/6119 | 5.0% | Insertion | Frameshift |
| TEVs-PEST | +1357 | G>T | 360/5983 | 6.0% | Substitution | Missense G453X |
| Intergenic | +1564 | +A | 946/6133 | 15.4% | Insertion | - |
| P | +1669 | +A | 138/6133 | 2.3% | Insertion | Frameshift |

| SIR-C-LowTEVp-P6 bc4—bc7 | | | | | | |
|---|---|---|---|---|---|---|
| | Position | Variant | N (q>20) | Freq % | Mutation | Effect on CDS |
| Upstream N | -49 | +A | 494/5209 | 9.5% | Insertion | - |
| | -20 | +G | 123/5209 | 2.4% | Insertion | - |
| | -19 | A>G | 2864/4984 | 5.7% | Substitution | - |
| N gene | +108 | +A | 167/5210 | 3.2% | Insertion | Frameshift |
| | +166 | +T | 136/5210 | 2.6% | Insertion | Frameshift |
| | +237 | +T | 400/5210 | 7.7% | Insertion | Frameshift |
| | +245 | +G | 123/5210 | 2.4% | Insertion | Frameshift |
| | +466 | +A | 146/5213 | 2.8% | Insertion | Frameshift |
| | +636 | +T | 261/5214 | 5.0% | Insertion | Frameshift |
| TEVs-PEST | +1357 | G>T | 546/5066 | 10.8% | Substitution | Missense G453X |
| Intergenic | +1564 | +A | 816/5212 | 15.7% | Insertion | - |
| P | +1669 | +A | 120/5212 | 2.3% | Insertion | Frameshift |

| SIR-D-LowTEVp-P6 bc4—bc7 | | | | | | |
|---|---|---|---|---|---|---|
| | Position | Variant | N (q>20) | Freq % | Mutation | Effect on CDS |
| Upstream N | -49 | +A | 492/5279 | 9.3% | Insertion | - |
| | -21 | -N | 114/5279 | 2.2% | Deletion | - |
| | -20 | +G | 119/5279 | 2.3% | Insertion | - |
| | -19 | A>G | 1553/5049 | 30.8% | Substitution | - |
| | -9 | A>T | 104/5189 | 2.0% | Substitution | - |
| N gene | +108 | +A | 163/5279 | 3.1% | Insertion | Frameshift |
| | +166 | +T | 129/5279 | 2.4% | Insertion | Frameshift |
| | +237 | +T | 434/5279 | 8.2% | Insertion | Frameshift |
| | +245 | +G | 106/5279 | 2.0% | Insertion | Frameshift |
| | +466 | +A | 148/5281 | 2.8% | Insertion | Frameshift |
| | +612 | +T | 120/5281 | 2.3% | Insertion | Frameshift |
| | +636 | +T | 279/5281 | 5.3% | Insertion | Frameshift |
| TEVs-PEST | +1357 | - | - | - | - | - |
| Intergenic | +1564 | +A | 831/5281 | 15.7% | Insertion | - |
| P | +1669 | +A | 123/5281 | 2.3% | Insertion | Frameshift |

| SIR-A-HighTEVp-P8 bc6—bc7 | | | | | | |
|---|---|---|---|---|---|---|
| | Position | Variant | N (q>20) | Freq % | Mutation | Effect on CDS |

| | | | | | | |
|---|---|---|---|---|---|---|
| Upstream N | -49 | +A | 541/6868 | 7.9% | Insertion | - |
| | -21 | -N | 299/6868 | 4.4% | Deletion | - |
| | -20 | +G | 431/6868 | 6.3% | Insertion | - |
| | -19 | A>G | 3684/6150 | 60.0% | Substitution | - |
| N gene | +108 | +A | 198/6867 | 2.9% | Insertion | Frameshift |
| | +166 | +T | 157/6867 | 2.3% | Insertion | Frameshift |
| | +237 | +T | 583/6867 | 8.5% | Insertion | Frameshift |
| | +245 | +G | 138/6867 | 2.0% | Insertion | Frameshift |
| | +466 | +A | 181/6868 | 2.6% | Insertion | Frameshift |
| | +636 | +T | 342/6870 | 5.0% | Insertion | Frameshift |
| TEVs-PEST | +1357 | G>T | 651/6620 | 9.8% | Substitution | Missense G453X |
| Intergenic | +1564 | +A | 952/6896 | 13.8% | Insertion | - |
| P | +1669 | +A | 144/6898 | 2.1% | Insertion | Frameshift |

| SIR-B-HighTEVp-P8 bc6—bc8 | | | | | | |
|---|---|---|---|---|---|---|
| | Position | Variant | N (q>20) | Freq % | Mutation | Effect on CDS |
| Upstream N | -49 | +A | 571/6246 | 9.1% | Insertion | - |
| | -21 | -N | 182/6246 | 2.9% | Deletion | - |
| | -20 | +G | 319/6246 | 5.1% | Insertion | - |
| | -19 | A>G | 3836/5763 | 66.6% | Substitution | - |
| | -18 | A>C | 171/5940 | 2.9% | Substitution | - |
| N gene | +108 | +A | 197/6247 | 3.2% | Insertion | Frameshift |
| | +166 | +T | 167/6247 | 2.7% | Insertion | Frameshift |
| | +237 | +T | 486/6247 | 7.8% | Insertion | Frameshift |
| | +245 | +G | 145/6248 | 2.3% | Insertion | Frameshift |
| | +466 | +A | 149/6249 | 2.4% | Insertion | Frameshift |
| | +636 | +T | 323/6251 | 5.2% | Insertion | Frameshift |
| TEVs-PEST | +1357 | G>T | 365/6068 | 6.0% | Substitution | Missense G453X |
| Intergenic | +1564 | +A | 927/6259 | 14.8% | Insertion | - |
| P | +1669 | +A | 152/6259 | 2.4% | Insertion | Frameshift |

| SIR-C-HighTEVp-P8 bc6—bc9 | | | | | | |
|---|---|---|---|---|---|---|
| | Position | Variant | N (q>20) | Freq % | Mutation | Effect on CDS |
| Upstream N | -49 | +A | 598/6403 | 9.3% | Insertion | - |
| | -19 | A>G | 6024/6304 | 95.6% | Substitution | - |
| N gene | +108 | +A | 200/6404 | 3.1% | Insertion | Frameshift |
| | +166 | +T | 146/6404 | 2.3% | Insertion | Frameshift |
| | +237 | +T | 518/6405 | 8.1% | Insertion | Frameshift |
| | +245 | +G | 158/6405 | 2.5% | Insertion | Frameshift |
| | +466 | +A | 172/6406 | 2.7% | Insertion | Frameshift |
| | +636 | +T | 311/6407 | 4.9% | Insertion | Frameshift |
| TEVs-PEST | - | - | - | - | - | - |
| Intergenic | +1564 | +A | 986/6410 | 15.4% | Insertion | - |
| P | +1669 | +A | 139/6408 | 2.2% | Insertion | Frameshift |

| SIR-D-HighTEVp-P8 bc6—bc10 | | | | | | |
|---|---|---|---|---|---|---|
| | Position | Variant | N (q>20) | Freq % | Mutation | Effect on CDS |
| Upstream N | -49 | +A | 482/5760 | 8.4% | Insertion | - |
| | -19 | A>G | 5092/5625 | 9.1% | Substitution | - |
| | -18 | A>G | 155/5609 | 2.8% | Substitution | - |
| | -9 | A>T | 247/5402 | 4.6% | Substitution | - |
| | -9 | A>G | 449/5402 | 8.3% | Substitution | - |
| | -9 | +G | 120/5761 | 2.1% | Insertion | - |
| | -6 | C>T | 680/5586 | 12.2% | Substitution | - |
| | -6 | +T | 167/5761 | 2.9% | Insertion | - |
| | -5 | C>A | 153/5412 | 2.8% | Substitution | - |
| N gene | +108 | +A | 163/5763 | 2.8% | Insertion | Frameshift |
| | +166 | +T | 119/5763 | 2.1% | Insertion | Frameshift |
| | +237 | +T | 414/5763 | 7.2% | Insertion | Frameshift |
| | +466 | +A | 119/5764 | 2.1% | Insertion | Frameshift |
| | +612 | +T | 127/5764 | 2.2% | Insertion | Frameshift |
| | +636 | +T | 291/5764 | 5.0% | Insertion | Frameshift |
| TEVs-PEST | - | - | - | - | - | - |
| Intergenic | +1564 | +A | 861/5766 | 14.9% | Insertion | - |
| P | +1669 | +A | 137/5766 | 2.4% | Insertion | Frameshift |

| SIR-A-LowTEVp-P8 bc4—bc9 | | | | | | |
|---|---|---|---|---|---|---|
| | Position | Variant | N (q>20) | Freq % | Mutation | Effect on CDS |
| Upstream N | -49 | +A | 646/7058 | 9.2% | Insertion | - |
| | -21 | -N | 252/7059 | 3.6% | Deletion | - |
| | -20 | +G | 417/7059 | 5.9% | Insertion | - |
| | -19 | A>G | 2752/6358 | 43.3% | Substitution | - |
| | -6 | C>T | 171/6942 | 2.5% | Substitution | - |
| | -5 | C>A | 542/6530 | 8.3% | Substitution | - |
| N gene | +108 | +A | 346/7058 | 4.9% | Insertion | Frameshift |
| | +166 | +T | 178/7058 | 2.5% | Insertion | Frameshift |
| | +237 | +T | 622/7058 | 8.8% | Insertion | Frameshift |
| | +245 | +G | 161/7058 | 2.3% | Insertion | Frameshift |
| | +466 | +A | 194/7059 | 2.7% | Insertion | Frameshift |
| | +612 | +T | 150/7060 | 2.1% | Insertion | Frameshift |
| | +636 | +T | 345/7060 | 4.9% | Insertion | Frameshift |
| | +795 | T>C | 1604/6265 | 25.6% | Substitution | Silent F265 |
| | +795 | +C | 318/7061 | 4.5% | Insertion | Frameshift |
| TEVs-PEST | +1357 | G>T | 1122/6684 | 16.8% | Substitution | Missense G453X |
| Intergenic | +1564 | +A | 1079/7085 | 15.2% | Insertion | - |
| P | +1669 | +A | 161/7090 | 2.3% | Insertion | Frameshift |

| SIR-B-LowTEVp-P8 bc4—bc10 | | | | | | |
|---|---|---|---|---|---|---|
| | Position | Variant | N (q>20) | Freq % | Mutation | Effect on CDS |
| Upstream N | -49 | +A | 647/6759 | 9.6% | Insertion | - |

| | | | | | | |
|---|---|---|---|---|---|---|
| | -21 | -N | 242/6761 | 3.6% | Deletion | - |
| | -20 | +G | 371/6761 | 5.5% | Insertion | - |
| | -19 | A>G | 2200/6168 | 35.7% | Substitution | - |
| | -18 | A>C | 400/6309 | 6.3% | Substitution | - |
| N gene | +108 | +A | 224/6761 | 3.3% | Insertion | Frameshift |
| | +166 | +T | 157/6761 | 2.3% | Insertion | Frameshift |
| | +237 | +T | 575/6760 | 8.5% | Insertion | Frameshift |
| | +466 | +A | 189/6764 | 2.8% | Insertion | Frameshift |
| | +636 | +T | 353/6763 | 5.2% | Insertion | Frameshift |
| | +1349 | C>A | 144/6671 | 2.2% | Substitution | Missense S450X |
| TEVs-PEST | +1357 | G>T | 1192/6372 | 18.7% | Substitution | Missense G453X |
| Intergenic | +1564 | +A | 1026/6769 | 15.2% | Insertion | - |
| P | +1669 | +A | 173/6772 | 2.6% | Insertion | Frameshift |

| SIR-C-LowTEVp-P8 bc4—bc11 | | | | | | |
|---|---|---|---|---|---|---|
| | Position | Variant | N (q>20) | Freq % | Mutation | Effect on CDS |
| Upstream N | -49 | +A | 614/6893 | 8.9% | Insertion | - |
| | -20 | +G | 261/6893 | 3.8% | Insertion | - |
| | -19 | A>G | 5317/6466 | 82.2% | Substitution | - |
| N gene | +108 | +A | 215/6894 | 3.1% | Insertion | Frameshift |
| | +237 | +T | 564/6894 | 8.2% | Insertion | Frameshift |
| | +466 | +A | 207/6895 | 3.0% | Insertion | Frameshift |
| | +636 | +T | 364/6895 | 5.3% | Insertion | Frameshift |
| TEVs-PEST | +1357 | G>T | 1013/6551 | 15.5% | Substitution | Missense G453X |
| Intergenic | +1564 | +A | 1053/6920 | 15.2% | Insertion | - |
| P | - | - | - | - | - | - |

| SIR-D-LowTEVp-P8 bc4—bc12 | | | | | | |
|---|---|---|---|---|---|---|
| | Position | Variant | N (q>20) | Freq % | Mutation | Effect on CDS |
| Upstream N | -49 | +A | 541/5872 | 9.2% | Insertion | - |
| | -20 | +G | 190/5872 | 3.2% | Insertion | - |
| | -19 | A>G | 4259/5565 | 76.5% | Substitution | - |
| | -9 | A>T | 141/5738 | 2.5% | Substitution | - |
| N gene | +108 | +A | 168/5876 | 2.9% | Insertion | Frameshift |
| | +166 | +T | 154/5876 | 2.6% | Insertion | Frameshift |
| | +237 | +T | 491/5876 | 8.4% | Insertion | Frameshift |
| | +245 | +G | 133/5876 | 2.3% | Insertion | Frameshift |
| | +332 | +A | 123/5876 | 2.1% | Insertion | Frameshift |
| | +466 | +A | 152/5876 | 2.6% | Insertion | Frameshift |
| | +612 | +T | 134/5876 | 2.3% | Insertion | Frameshift |
| | +636 | +T | 324/5876 | 5.5% | Insertion | Frameshift |
| TEVs-PEST | +1357 | G>T | 521/5707 | 9.1% | Substitution | Missense G453X |
| Intergenic | +1564 | +A | 996/5881 | 17.0% | Insertion | - |
| P | +1669 | +A | 150/5882 | 2.6% | Insertion | Frameshift |

**Table 2**

List of detected mutations above 2% thresholds in SiR viruses amplified in high- and low-TEVp packaging cells sequenced by SMRT NGS sequencing. The position of the mutations is defined considering +1 the first base of the nucleoprotein N coding sequence.
