## Supplementary material for "Genomic stability of Self-inactivating Rabies": Table 3

### NGS sequencing results of purified viruses used *in vivo*

| SIR-CRE purified bc3—bc5 | | | | | | |
|---|---|---|---|---|---|---|
| | Position | Variant | N (q>20) | Freq % | Mutation | Effect on CDS |
| Upstream N | -49 | +A | 238/5196 | 4.6% | Insertion | - |
| N gene | +237 | +T | 199/5196 | 3.8% | Insertion | Frameshift |
| | +636 | +T | 150/5200 | 2.9% | Insertion | Frameshift |
| TEVs-PEST | - | - | - | - | - | - |
| Intergenic | +1564 | +A | 544/5205 | 10.5% | Insertion | - |
| P | - | - | - | - | - | - |

| SIR-CRE purified, 1 week p.i. *in vivo* (A) bc5—bc10 | | | | | | |
|---|---|---|---|---|---|---|
| | Position | Variant | N (q>20) | Freq % | Mutation | Effect on CDS |
| Upstream N | -49 | +A | 474/5211 | 9.1% | Insertion | - |
| | -21 | +A | 110/5211 | 2.1% | Insertion | - |
| N gene | +108 | +A | 176/5211 | 3.4% | Insertion | Frameshift |
| | +166 | +T | 132/5211 | 2.5% | Insertion | Frameshift |
| | +237 | +T | 389/5211 | 7.5% | Insertion | Frameshift |
| | +245 | +G | 108/5211 | 2.1% | Insertion | Frameshift |
| | +466 | +A | 135/5211 | 2.6% | Insertion | Frameshift |
| | +612 | +T | 108/5210 | 2.1% | Insertion | Frameshift |
| | +636 | +T | 288/5210 | 5.5% | Insertion | Frameshift |
| TEVs-PEST | - | - | - | - | - | - |
| Intergenic | +1564 | +A | 773/5213 | 14.8% | Insertion | - |
| P | +1669 | +A | 128/5213 | 2.5% | Insertion | Frameshift |

| SIR-CRE purified, 1 week p.i. *in vivo* (B) bc5—bc11 | | | | | | |
|---|---|---|---|---|---|---|
| | Position | Variant | N (q>20) | Freq % | Mutation | Effect on CDS |
| Upstream N | -49 | +A | 482/5542 | 8.7% | Insertion | - |
| N gene | +108 | +A | 157/5543 | 2.8% | Insertion | Frameshift |
| | +166 | +T | 125/5543 | 2.3% | Insertion | Frameshift |
| | +237 | +T | 402/5543 | 7.3% | Insertion | Frameshift |
| | +245 | +G | 123/5543 | 2.2% | Insertion | Frameshift |
| | +466 | +A | 157/5543 | 2.8% | Insertion | Frameshift |
| | +612 | +T | 112/5543 | 2.0% | Insertion | Frameshift |
| | +636 | +T | 276/5543 | 5.0% | Insertion | Frameshift |
| TEVs-PEST | - | - | - | - | - | - |
| Intergenic | +1564 | +A | 744/5542 | 13.4% | Insertion | - |
| P | +1669 | +A | 144/5542 | 2.6% | Insertion | Frameshift |

| SIR-CRE purified, 1 week p.i. *in vivo* (C) bc5—bc12 | | | | | | |
|---|---|---|---|---|---|---|
| | Position | Variant | N (q>20) | Freq % | Mutation | Effect on CDS |
| Upstream N | -49 | +A | 481/5150 | 9.3% | Insertion | - |
| N gene | +108 | +A | 137/5150 | 2.7% | Insertion | Frameshift |
| | +166 | +T | 118/5150 | 2.3% | Insertion | Frameshift |
| | +237 | +T | 390/5150 | 7.6% | Insertion | Frameshift |
| | +245 | +G | 104/5150 | 2.0% | Insertion | Frameshift |

| | | | | | | |
|---|---|---|---|---|---|---|
| | +466 | +A | 140/5150 | 2.7% | Insertion | Frameshift |
| | +612 | +T | 116/5150 | 2.3% | Insertion | Frameshift |
| | +636 | +T | 255/5150 | 5.0% | Insertion | Frameshift |
| TEVs-PEST | - | - | - | - | - | - |
| Intergenic | +1564 | +A | 739/5148 | 14.4% | Insertion | - |
| P | +1669 | +A | 130/5148 | 2.5% | Insertion | Frameshift |

| SIR-G453X-CRE purified bc3—bc11 | | | | | | |
|---|---|---|---|---|---|---|
| | Position | Variant | N (q>20) | Freq % | Mutation | Effect on CDS |
| Upstream N | -49 | +A | 211/4886 | 4.3% | Insertion | - |
| N gene | +237 | +T | 244/4890 | 5.0% | Insertion | Frameshift |
| | +636 | +T | 138/4911 | 2.8% | Insertion | Frameshift |
| TEVs-PEST | +1357 | G>T | 4780/4912 | 97.3% | Substitution | Missense G453X |
| Intergenic | +1564 | +A | 502/4924 | 10.2% | Insertion | - |
| P | - | - | - | - | - | - |

**Table 3**

List of detected mutations above 2% threshold in SiR viruses amplified in high- and low-TEVp packaging cells sequenced by SMRT NGS sequencing. The position of the mutations is defined considering +1 the first base of the nucleoprotein N coding sequence.
